# Mitochondrial CaMKII causes metabolic reprogramming, energetic insufficiency, and dilated cardiomyopathy

**DOI:** 10.1101/2020.02.13.947564

**Authors:** Elizabeth D. Luczak, Yuejin Wu, Jonathan M. Granger, Mei-ling A. Joiner, Nicholas R. Wilson, Ashish Gupta, Priya Umapathi, Kevin R. Murphy, Oscar E. Reyes Gaido, Amin Sabet, Eleonora Corradini, Yibin Wang, Albert J.R. Heck, An-Chi Wei, Robert G. Weiss, Mark E. Anderson

## Abstract

Despite the clear association between myocardial injury, heart failure and depressed myocardial energetics, little is known about upstream signals responsible for remodeling myocardial metabolism after pathological stress. We found increased mitochondrial calmodulin kinase II (CaMKII) activation and left ventricular dilation in mice one week after myocardial infarction (MI) surgery. In contrast, mice with genetic mitochondrial CaMKII inhibition were protected from left ventricular dilation and dysfunction after MI. Mice with myocardial and mitochondrial CaMKII over-expression (mtCaMKII) had severe dilated cardiomyopathy and decreased ATP that caused elevated cytoplasmic resting (diastolic) Ca^2+^ concentration and reduced mechanical performance. We mapped a metabolic pathway that allowed us to rescue disease phenotypes in mtCaMKII mice, providing new insights into physiological and pathological metabolic consequences of CaMKII signaling in mitochondria. Our findings suggest myocardial dilation, a disease phenotype lacking specific therapies, can be prevented by targeted replacement of mitochondrial creatine kinase, or mitochondrial-targeted CaMKII inhibition.

## Introduction

Heart failure is a leading cause of death worldwide, and failing myocardium is marked by energetic defects ^1, 2^. Loss of creatine kinase ^3^ and decreased mitochondrial respiration ^4^ are common findings in myocardium after pathological injury, including myocardial infarction (MI), a major cause of heart failure. Although failing hearts have reduced energy reserves, little is known about upstream control points contributing to decreased energy metabolism, nor the specific structural and functional consequences of these defects on myocardial responses to injury. MI triggers a complex set of changes in myocardium, including cardiomyocyte death and cardiac fibrosis, hypertrophy, and chamber dilation. Collectively these changes are termed adverse structural remodeling. Much is known about cellular signals contributing to myocardial hypertrophy, fibrosis, and myocyte death ^5, 6, 7^, whereas relatively little is known about molecular signals causing myocardial dilation ^8^. Although hypertrophy has been proposed to be a precondition for myocardial chamber dilation ^9^, the connection between myocardial hypertrophy and cardiac chamber dilation is uncertain. The nature of the relationship between myocardial hypertrophy and cardiac chamber dilation is important clinically because current therapies are inadequate for treating or preventing pathological cardiac chamber dilation. Emergent evidence indicates that defects in mitochondrial function and energy depletion may contribute to heart failure and dilated cardiomyopathy ^10^.

The multifunctional Ca^2+^ and calmodulin dependent protein kinase II (CaMKII) is a pluripotent signal for promoting cardiomyopathy ^11^. Mouse models of myocardial CaMKII over-expression, without subcellular targeting, show activation of myocardial hypertrophy gene programs, increased cell death, fibrosis, chamber dilation, and premature death ^12, 13^. Excessive CaMKII activity causes myocardial hypertrophy by catalyzing phosphorylation of HDAC4, leading to cytoplasmic partitioning of HDAC4, and derepression of hypertrophic transcriptional programs ^14, 15^. CaMKII promotes myocardial death via multiple pathways ^16^, including induction of pro-apoptotic genes through NF-κB activation ^17, 18^, defective DNA repair^19^, promoting cytosolic Ca^2+^ overload ^20^, and triggering mitochondrial permeability transition pore opening ^21^. CaMKII also promotes dilated cardiomyopathy ^13, 22^, but the mechanism is unknown. CaMKII is present in the mitochondrial fraction of heart and vascular lysates ^23, 24^, and is found in the mitochondrial matrix ^25^, but responses to sustained, pathological mitochondrial activation in heart are unknown.

Here we identified increased mitochondrial CaMKII activation in failing mouse hearts one week after myocardial infarction surgery, and found that mouse hearts with myocardial and mitochondrial matrix CaMKII inhibition, due to transgenic expression of a potent, and selective CaMKII inhibitor polypeptide (mtCaMKIIN) ^25^, were protected from left ventricular dilation and dysfunction one week after myocardial infarction. We developed a new genetic mouse model with myocardial- and mitochondrial-targeted CaMKII over-expression, and found that excessive mitochondrial CaMKII causes dilated cardiomyopathy, without myocardial hypertrophy or death, by reducing expression of assembled complex I and the mitochondrial isoform of creatine kinase (CKmito). Genetic replacement of CKmito but not cytosolic myofibrillar creatine kinase (CK-M) was sufficient to rescue myocardial energetics, restore intracellular Ca^2+^ homeostasis, and prevent dilated cardiomyopathy. Mitochondrial CaMKII over-expression coherently affected TCA cycle dehydrogenases, causing increased phosphorylation, activity and heightened production of NADH, an essential electron donor for oxidative phosphorylation. These findings show excessive mitochondrial CaMKII can trigger dilated cardiomyopathy, independent of myocardial hypertrophy and cell death, by depressing energetics. Our findings suggest that mitochondrial CaMKII selectively contributes to cardiac chamber dilation, potentially in acquired forms of myocardial injury with high importance to public health, and that dilated cardiomyopathy can be prevented by therapies capable of restoring metabolism.

## Results

### Increased mitochondrial CaMKII activity in failing mouse heart

Myocardial infarction is a major cause of heart failure, one of the largest, unsolved public health challenges ^26, 27^. Myocardial CaMKII activity is increased in myocardial lysates from heart failure patients, and in many animal models of heart failure ^28^. Increased CaMKII is known to contribute to myocardial hypertrophy and arrhythmias by phosphorylation of nuclear and cytoplasmic target proteins ^29, 30, 31^. CaMKII was recently identified in mitochondria ^23, 24, 25^, and mitochondrial CaMKII inhibition protects against myocardial death, acutely, during the first hours after myocardial infarction ^25^. However, the potential for excessive, chronic, mitochondrial CaMKII to contribute to specific myocardial disease phenotypes is unexplored. As a first step, we tested whether mitochondrial CaMKII activity is increased in failing hearts by measuring total and threonine 287 auto-phosphorylated CaMKII, a marker of CaMKII activation ^32^, in mitochondria isolated from mouse hearts following myocardial infarction or sham surgery. We found an increase in auto-phosphorylated CaMKII in mitochondria isolated from heart tissue one week after myocardial infarction surgery, compared to sham operated controls (Fig 1A and B). The total amount of CaMKII in the mitochondria was similar between the myocardial infarcted and sham hearts. We also observed an increase in phosphorylated CaMKII in the cytosol from the same hearts (Fig S1A). Additionally, we quantified phosphorylation of a known mitochondrial target of CaMKII, the mitochondrial Ca^2+^ uniporter (MCU) ^24, 25^. Rapid mitochondrial matrix Ca^2+^ entry is mediated by MCU ^33, 34^, and CaMKII increases MCU Ca^2+^ entry ^24, 25^, although this finding has been disputed ^35, 36, 37^. We used recently validated phospho-specific antibodies against MCU (pMCU) ^24^, predicted CaMKII phosphorylation sites, and found an increase in phosphorylation of this CaMKII target in mitochondria from hearts after MI compared to sham operated hearts (Fig 1A and B). Collectively, these results suggest that increased mitochondrial CaMKII activity occurs in response to myocardial infarction, and persists substantially beyond the perioperative episode.

**Figure 1.**
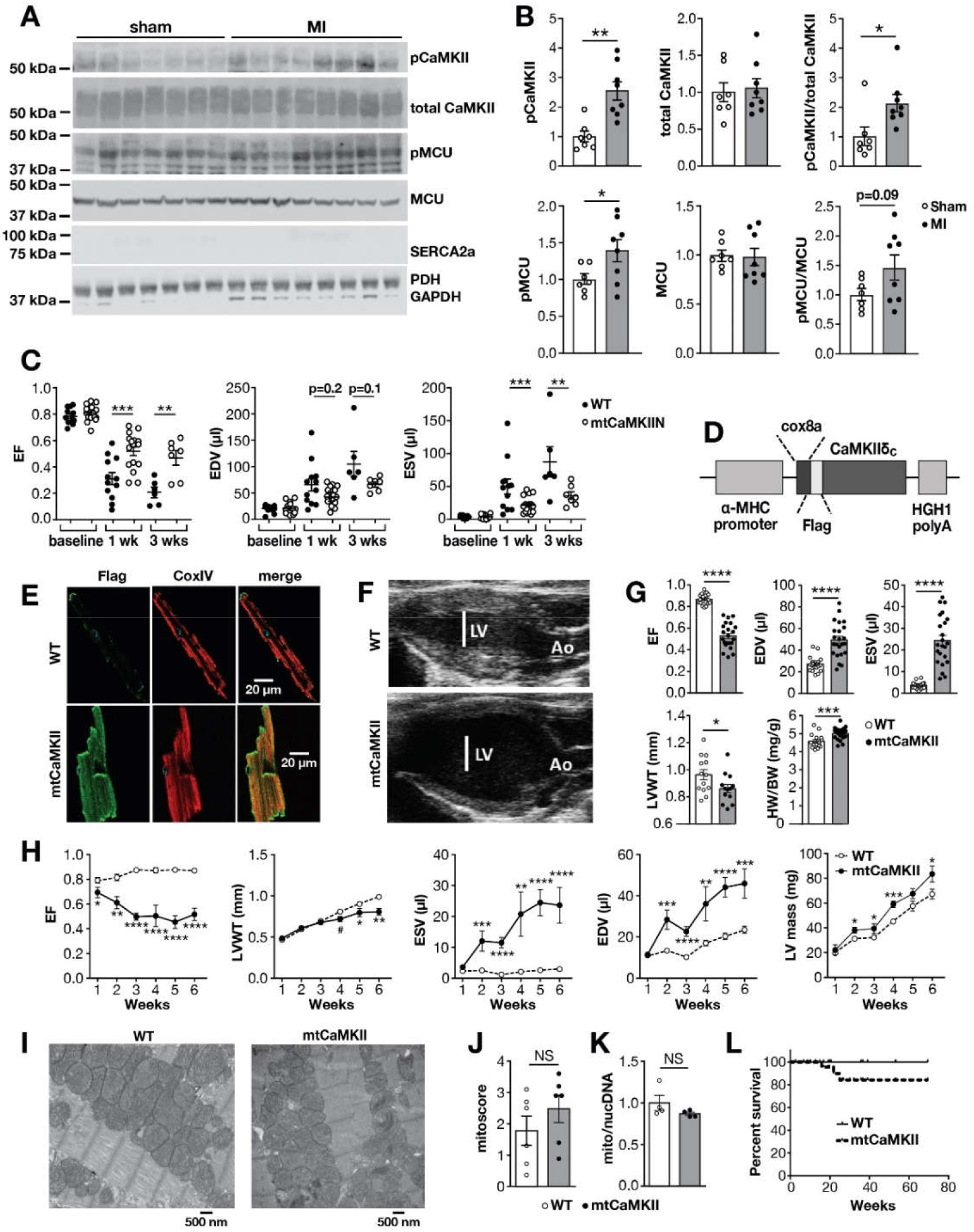
Mitochondrial CaMKII is activated by myocardial infarction and causes dilated cardiomyopathy. (A) Western blot of mitochondrial lysates and (B) summary data for phosphorylated CaMKII and total CaMKII and phosphorylated MCU and total MCU normalized to coomassie staining for sham (n=7) and MI (n=8) mitochondria. Blots for cellular compartment markers also included: SERCA2a (SR membrane), PDH (mitochondrial matrix), GAPDH (cytosol). (C) Summary data from echocardiographic measurements from WT (n=6-12) and mtCaMKIIN (n=7-16) mice before, 1 and 3 weeks after MI. EF (left ventricular ejection fraction), EDV (left ventricular end diastolic volume), ESV (left ventricular end systolic volume) (D) Schematic of the mtCaMKII transgene construct, α-MHC: α-myosin heavy chain, cox8a: cox8a mitochondrial localization sequence, Flag: flag epitope tag, HGH1 polyA: human growth hormone polyA signal. (E) Immunofluorescence micrographs of isolated ventricular myocytes from WT and mtCaMKII mice (F) Representative echocardiographic images in end diastole, LV: left ventricle, Ao: aorta (G) Summary data for echocardiographic measurements from adult WT (n=13-20) and mtCaMKII (n=13-23) mice. LVWT (left ventricular wall thickness), HW/BW (heart weight/body weight) (H) Summary data for echocardiographic measurements from young (1-6 weeks of age) WT (n=12) and mtCaMKII (n=6) mice. (I) Representative transmission electron micrographs from WT and mtCaMKII left ventricle. (J) Mitochondrial injury score for WT (n=6) and mtCaMKII (n=6) hearts. (K) Mitochondrial DNA content (cox1) normalized to nuclear DNA (βglobin) for WT (n=4) and mtCaMKII (n=4) hearts. (L) Kaplan-Meier survival relationship for WT (n=41) and mtCaMKII (n=48) mice. (Data are represented as mean ± SEM, significance was determined using a two-tailed t test or log-rank test (survivorship), ****P<0.0001, ***P<0.001, **P<0.01, *P<0.05, NS=not significant)

### Mitochondrial CaMKII inhibition protects against left ventricular dilation after MI

Given the increased mitochondrial CaMKII activity in mice after myocardial infarction, we asked whether inhibition of mitochondrial CaMKII, by myocardial targeted over-expression of CaMKIIN (mtCaMKIIN) ^25^, could protect the heart from chronic responses to injury. We next measured mtCaMKIIN expression and found it was present in mitochondrial and extramitochondrial fractions (Fig S1B). CaMKIIN is expressed endogenously in neurons, but not in heart ^38^, and mtCaMKIIN hearts are protected against mitochondrial Ca^2+^ overload, and increased myocardial death in the first hours after pathological stress ^25^. Here we subjected mtCaMKIIN mice and WT littermates to surgical MI, and measured heart size and function using echocardiography over three weeks. Although basal myocardial function, prior to MI surgery (Fig 1C), and infarcted myocardial area 24 hours after myocardial infarction surgery (Fig S1C) were similar between mtCaMKIIN and WT littermate controls, the WT mice showed worse left ventricular ejection fractions, and progressed to more severe left ventricular dilatation than the mtCaMKIIN mice by one week after myocardial infarction. The differences in left ventricular dilation and function between WT and mtCaMKIIN hearts were sustained for three weeks after surgery, suggesting mitochondrial CaMKII participated in structural remodeling on a subacute or chronic timescale (Fig 1C). In order to verify that mitochondrial CaMKII activity was decreased in the mtCaMKIIN hearts, we quantified pMCU and found a trend for reduced phosphorylation of this CaMKII target 1 week following MI (Fig S1D). These data further support the role of mitochondrial CaMKII in structural remodeling following cardiac injury.

CaMKII has been shown to contribute to cardiac inflammation following ischemic injury ^17, 18, 39^. In order to determine if inhibition of mitochondrial CaMKII reduces inflammation after MI, we measured chemokine expression and leukocyte infiltration 1 week after coronary ligation. We did find evidence of increased inflammation after MI surgery, but found no difference in CCL2 or CCL3 mRNA expression (Fig S1E), nor CD45+ cell infiltration (Fig S1F) in mtCaMKII hearts compared to WT littermates. These data suggest that inhibition of mitochondrial CaMKII does not affect the inflammatory response in this model of injury.

### Mitochondrial CaMKII over-expression causes dilated cardiomyopathy

To directly test the effects of increased CaMKII activity in cardiac mitochondria, we developed a mouse model with chronic transgenic over-expression of CaMKII targeted to myocardial mitochondrial matrix (mtCaMKII). We fused the CaMKIIδ cDNA to a *Cox8a* mitochondrial localization sequence and a flag epitope tag, and expressed this construct under the control of the α myosin heavy chain promotor (Fig 1D) ^40^. Cardiac myocytes isolated from mtCaMKII mice showed flag-CaMKII localized primarily with a mitochondrial marker (CoxIV) (Fig 1E). The mtCaMKII mice developed dilated cardiomyopathy with reduced left ventricular ejection fraction, increased left ventricular volumes, and decreased left ventricular wall thickness (Fig 1F and G). We observed modest cardiac hypertrophy in the mtCaMKII mice compared to WT littermates, as measured by heart weight normalized to body weight (Fig 1G, non-normalized data shown in Fig S2A), but cardiac myocyte cross-sectional area was not different in mtCaMKII hearts compared to WT littermate hearts (Fig S2B and C). Left ventricular dilation was apparent by two weeks of age, and preceded detectable hypertrophy in mtCaMKII hearts, compared to WT littermate controls (Fig 1H). Mitochondrial ultrastructural integrity scoring (Fig 1I and J) ^25^, and abundance, reflected by the ratio of mitochondrial to nuclear DNA (Fig 1K), were similar compared to WT littermate controls. These data show that mtCaMKII mice develop a severe dilated cardiomyopathy with minimal myocardial hypertrophy.

### Dilated cardiomyopathy persists in mtCaMKII mice after extramitochondrial CaMKII inhibition

In contrast to the dilated cardiomyopathy and near normal lifespan in mtCaMKII mice (Fig 1L), myocardial CaMKII over expression in the absence of subcellular targeting causes marked cardiac hypertrophy, left ventricular dilation, increased myocardial cell death, cardiac fibrosis, heart failure, and premature death ^12, 13^. The striking differences in the phenotypes of mtCaMKII, and mice with transgenic myocardial CaMKII over-expression lacking subcellular targeting appeared to confirm a specific role for mitochondrial CaMKII in dilated cardiomyopathy. However, while the Cox8a was effective in predominantly targeting CaMKII over-expression to myocardial mitochondria, we also detected small amounts of mtCaMKII in extramitochondrial compartments (Fig 2A). To determine if extramitochondrial mtCaMKII contributed to cardiomyopathy in mtCaMKII mice, we crossed the mtCaMKII mice with mice expressing AC3-I, a CaMKII inhibitory peptide that is fused to enhanced green fluorescent protein (GFP) ^41^, and excluded from mitochondria (Fig 2B). The double transgenic mice, combining mitochondrial targeted CaMKII over-expression and extramitochondrial CaMKII inhibition, showed persistent left ventricular dysfunction (see left ventricular ejection fraction) and dilation (see left ventricular end systolic and diastolic volumes), but loss of left ventricular hypertrophy (see heart weight/body weight), compared to the mtCaMKII mice (Fig 2C). The partial improvement of left ventricular ejection fraction in the mtCaMKII x AC3-I interbred mice suggested that mitochondrial and extramitochondrial CaMKII each have the potential to decrease myocardial function, but that mitochondrial CaMKII exclusively promotes dilated cardiomyopathy, and not myocardial hypertrophy. We next crossed the mtCaMKII mice with mtCaMKIIN mice ^25^. The coexpression of mtCaMKIIN reversed the dilated cardiomyopathy phenotype in the mtCaMKII mice (Fig 2D), without reducing mtCaMKII expression (Fig 2E and F), confirming that dilated cardiomyopathy is due to excessive mitochondrial CaMKII activity, and excluding the possibility that dilated cardiomyopathy is a non-specific consequence of mitochondrial targeted transgenic protein over-expression. Considered together with previously published information ^13^, these data suggest that the dilated phenotype of the mtCaMKII heart is due to increased CaMKII activity in mitochondria, while myocardial hypertrophy and premature death are predominantly related to extramitochondrial actions of CaMKII.

**Figure 2.**
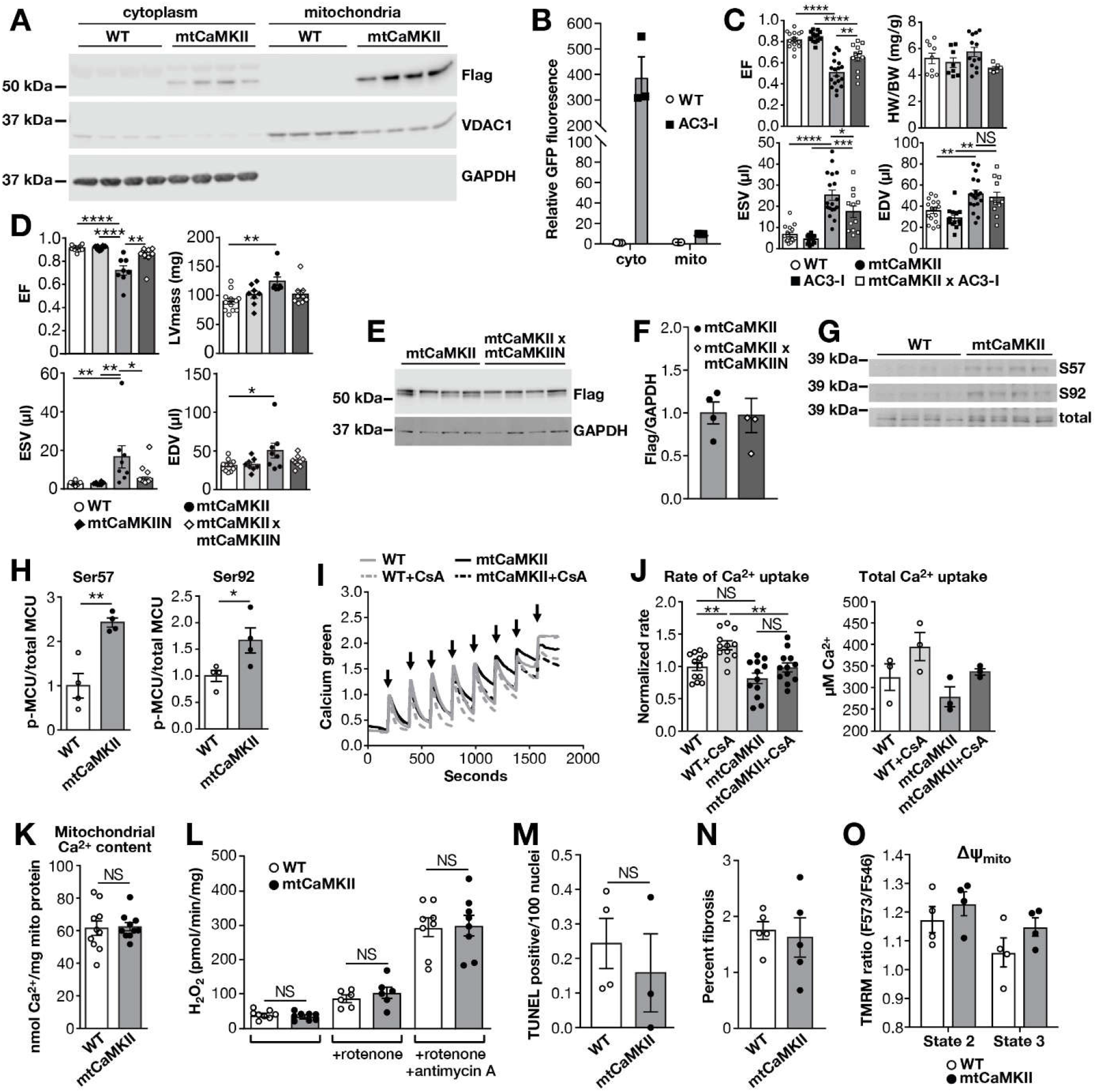
Dilated cardiomyopathy in mtCaMKII mice is independent of extramitochondrial CaMKII, MCU phosphorylation, mitochondrial Ca^2+^, ROS, and ΔΨ_mito_. (A) Western blot for flag-CaMKII, VDAC1 and GAPDH in cytoplasm and mitochondria fractions from WT (n=4) and mtCaMKII hearts (n=4) (B) Quantification AC3-I-GFP fusion protein expression in cytoplasm and mitochondria from WT and AC3-I mouse hearts (n=3). (C) Summary data for echocardiographic measurements from WT (n=16), AC3-I (n=13), mtCaMKII (n=18), and mtCaMKII x AC3-I (n=12) mice. (D) Summarized echocardiographic measurement data from WT (n=12), mtCaMKIIN (n=8), mtCaMKII (n=10), and mtCaMKII x mtCaMKIIN (n=10) mice. (E) Western blot and (F) summary data for flag-CaMKII normalized to GAPDH in mtCaMKII (n=4) and mtCaMKII x mtCaMKIIN (n=4) hearts. (G) Western blot and (H) summary data for phosphorylated MCU normalized to total MCU for WT (n=4) and mtCaMKII (n=4) mitochondria. (I) Mitochondrial Ca^2+^ uptake assay with cell membrane permeabilized adult ventricular myocytes from WT (n=3) and mtCaMKII (n=3) mice; arrows indicate addition of Ca^2+^. (J) Rate of Ca^2+^ uptake calculated from the first 3 peaks of Ca^2+^ addition and total Ca^2+^ uptake before mPTP opening. (K) Total mitochondrial Ca^2+^ content normalized to total protein in isolated mitochondria from WT (n=10) and mtCaMKII (n=10) hearts. (L) H2O2 production measured by Amplex Red in isolated mitochondria from WT (n=8) and mtCaMKII (n=8) hearts. (M) Quantification of TUNEL positive nuclei from heart sections of WT (5 sections from n=4 hearts) and mtCaMKII (5 sections from n=3 hearts) mice. (N) Quantification of fibrosis by Masson’s Trichrome staining from WT (6 sections from n=3 hearts) and mtCaMKII (6 sections from n=3 hearts) heart sections. (O) Mitochondrial membrane potential (ΔΨ _mito_) measured with TMRM under state 2 (substrate alone) and state 3 (substrate plus ADP) respiration in isolated mitochondria from WT (n=4) and mtCaMKII (n=4) hearts. (Data are represented as mean ± SEM, significance was determined using a two-tailed t test or 1 way AVOVA with Tukey’s multiple comparison’s test, ****P<0.0001, ***P<0.001, **P<0.01, *P<0.05, NS=not significant)

### No increased myocyte death and normal mitochondrial Ca^2+^, ROS, and ΔΨ_mito_ in mtCaMKII hearts

CaMKII plays a prominent role in myocardial biology, in part, by enhancing activity of intracellular Ca^2+^ homeostatic proteins ^11^. Mitochondrial targeted CaMKII inhibition, in mtCaMKIIN mice, protects against acute responses to myocardial injury by reducing cardiomyocyte death, Ca^2+^ overload, and loss of ΔΨ_mito_ ^25^. Mitochondria isolated from the hearts of mtCaMKII mice showed significantly increased pMCU (Fig 2G and H), suggesting excess phosphorylation at these sites in mtCaMKII compared to WT MCU. Based on these findings, we initially anticipated that dilated cardiomyopathy in mtCaMKII mice was related to mitochondrial Ca^2+^ overload, loss of ΔΨ_mito_, and increased myocardial death.

To test if mitochondrial Ca^2+^ entry was enhanced in mtCaMKII mitochondria, we measured MCU-mediated mitochondrial Ca^2+^ uptake, and mitochondrial Ca^2+^ content. We found that mitochondria from mtCaMKII and WT littermate control hearts had similar Ca^2+^ uptake rates (Fig 2I and J), and no differences in mitochondrial Ca^2+^ content (Fig 2K). In order to determine whether the mitochondrial transition pore contributed to the apparent rate of mitochondrial Ca^2+^ uptake, we also included cyclosporine A (CsA), an inhibitor of the mPTP, in the Ca^2+^ uptake assays (Fig 2I and J). We found an increase in the apparent rate of Ca^2+^ uptake in WT mitochondria, and a modest reduction in apparent rate of Ca^2+^ uptake in mtCaMKII compared to WT mitochondria in the presence of CsA, suggesting a reduction in mPTP activity in the mtCaMKII mitochondria. Taken together, these studies supported a view contrary to our starting hypothesis: that chronic mitochondrial CaMKII over-expression did not cause dilated cardiomyopathy by disrupting mitochondrial Ca^2+^ homeostasis. Furthermore, mtCaMKII mice did not exhibit increased mitochondrial ROS (Fig 2L), cardiomyocyte TUNEL staining (Fig 2M), or fibrosis (Fig 2N), and had similar ΔΨ_mito_ compared to WT littermate controls (Fig 2O). Measuring mitochondrial ROS is technically challenging ^42^, so we considered the possibility that our studies were inadequate to detect differences in ROS between WT and mtCaMKII hearts, potentially of a magnitude to contribute to the dilated cardiomyopathy in mtCaMKII mice. In order to more decisively determine if elevated ROS contributed to mtCaMKII cardiomyopathy, we took an orthogonal approach, and interbred mtCaMKII mice with mice transgenically over-expressing a mitochondrial targeted form of catalase (mCat), an enzyme that decomposes H_2_O_2_ into H_2_O and O_2_. The mCat mice are resistant to ROS mediated mitochondrial disease ^43^. The mtCaMKII x mCat interbred mice were not protected against left ventricular dilation, nor depressed ejection fraction compared to mtCaMKII mice (Fig S3A). Finally, *Nnt* is mutated in the C57BL/6J strain we used ^44^, suggesting that the left ventricular dilation in mtCaMKII cardiomyopathy phenotype may be artificially disconnected from ROS generation^45^. In order to examine this possibility, we back crossed mtCaMKII mice for 5 generations into the CD1 background that expresses WT *Nnt*. The mtCaMKII mice in the C57Bl/6J and CD1 genetic backgrounds exhibited similar left ventricular dilation (Fig S3B). Thus, in vitro and in vivo data strongly suggested that mtCaMKII cardiomyopathy was not a consequence of elevated ROS.

Given the lack of effect of increased mitochondrial CaMKII activity on myocyte survival, mitochondrial Ca^2+^, ROS or ΔΨ_mito_, activators and measures of the mitochondrial transition pore (mPTP) opening, we did not anticipate mPTP was involved in the dilated cardiomyopathy present in mtCaMKII hearts. However, to directly test for increased mPTP opening in mtCaMKII hearts in vivo we interbred mtCaMKII mice with mice lacking cyclophilin D (*Ppif*^−/−^), an mPTP subunit that is involved in pore opening ^46^. The *Ppif*^−/−^ mice are protected from mPTP opening ^47^. Consistent with our data up to this point, the *Ppif*^−/−^ x mtCaMKII interbred mice were not protected against dilated cardiomyopathy (Fig S3C). Taken together, these surprising in vitro and in vivo results strongly argued against defective mitochondrial Ca^2+^ or its consequences as a cause of cardiomyopathy in mtCaMKII mice.

### Reduced ATP production impairs cytoplasmic Ca^2+^ homeostasis in mtCaMKII hearts

Energy deficiency may contribute to dilated cardiomyopathy, ^48^, but little is known about specific upstream molecular signals initiating metabolic insufficiency. We initially found that the ATP concentration was significantly reduced in myocardial mitochondria lysates from mtCaMKII compared to WT littermate control hearts (Fig 3A). We next used ^31^P magnetic resonance (MR) spectroscopy to measure myocardial high-energy phosphates in vivo ^49, 50^. Cardiac ATP and creatine phosphate concentrations were significantly reduced, by 30%-45%, in mtCaMKII mice as compared to WT controls (Fig 3B and C). MR imaging confirmed dilated cardiomyopathy (Fig 3D, data supplement for cine), similar to our echocardiographic findings (Fig 1G). Together, these results show a significant reduction in ATP in mtCaMKII hearts, in vitro and in vivo, and are consistent with the hypothesis that impaired myocardial function in mtCaMKII mice is a consequence of energy deficiency.

**Figure 3.**
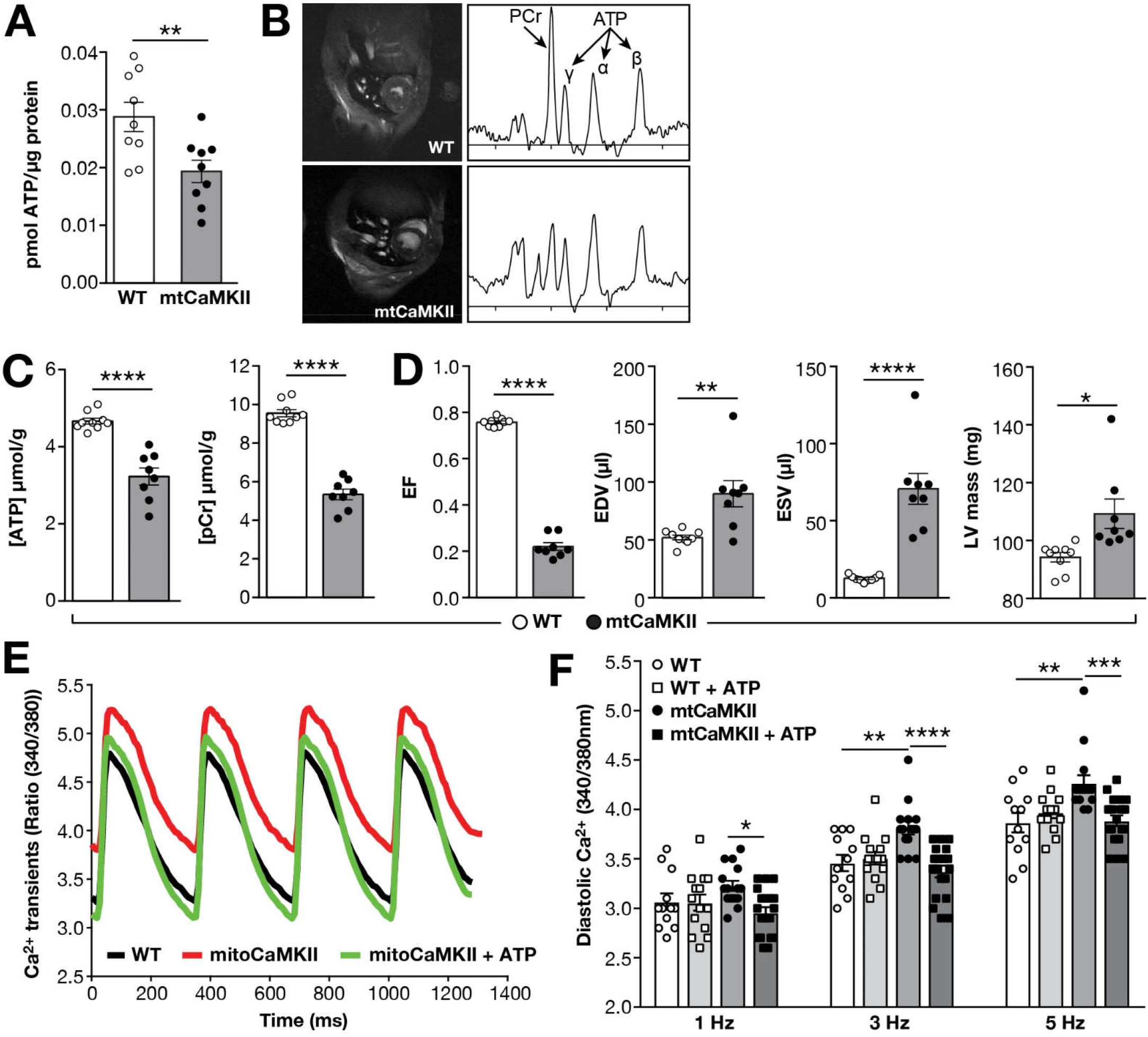
mtCaMKII hearts are ATP deficient. (A) ATP content in mitochondria isolated from WT (n=9) and mtCaMKII (n=9) hearts. (B) Example MRI images and corresponding ^31^P MR spectra and (C) summary data for ATP and creatine phosphate (PCr) quantification from in vivo ^31^P MR spectroscopy in WT (n=9) and mtCaMKII (n=8) hearts (D) Summary data from MRI measurements of WT and mtCaMKII hearts; see Fig S2 for MR Cines. (E) Example intracellular [Ca^2+^] signals recorded from Fura-2 and (F) summary data for diastolic (resting) [Ca^2+^] from isolated ventricular myocytes stimulated at 1, 3, or 5 Hz (n=12-19 ventricular myocytes isolated from 2-3 hearts/group). (Data are represented as mean ± SEM, significance was determined using a two-tailed t test or 1 way AVOVA with Tukey’s multiple comparison’s test, ****P<0.0001, ***P<0.001, **P<0.01, *P<0.05)

Cytoplasmic Ca^2+^ concentration ([Ca^2+^]_i_) controls cardiac contraction and relaxation, and [Ca^2+^]_i_ homeostasis represents a major physiological energy cost. We next measured [Ca^2+^]_i_ in ventricular myocytes isolated from mtCaMKII and WT hearts in response to increasing stimulation frequencies. Contrary to our expectations, these initial studies showed a decrease in resting [Ca^2+^]_i_ in cardiomyocytes isolated from mtCaMKII and WT hearts (Fig S4A) consistent with previous findings in hearts with CaMKII over-expression ^51^. We next considered that increased cytoplasmic CaMKII seen in mtCaMKII cardiomyocytes (Fig 2A) could mask effects of ATP deficiency on [Ca^2+^]_i_ in isolated, mechanically unloaded cardiomyocytes, potentially because energy requirements were too low to reveal differences. In order to test this idea, we repeated the [Ca^2+^]_i_ measurements in isolated ventricular myocytes dialyzed with AIP, a CaMKII inhibitory peptide. The AIP peptide was confined to the cytoplasmic compartment because it lacked a membrane permeation epitope and was dialyzed across the cell membrane using a microelectrode in whole cell mode patch clamp configuration. AIP dialysis had no effect on resting diastolic [Ca^2+^]_i_ in WT cardiomyocytes, but resulted in a significant, stimulation frequency dependent elevation in diastolic [Ca^2+^]_i_ (Fig 3E and F), unchanged peak Ca^2+^ amplitude (Fig S4B) and slower Ca^2+^ decay (Fig S4C) in mtCaMKII cardiomyocytes, consistent with the hypothesis that incompletely targeted mtCaMKII preserved cytoplasmic [Ca^2+^] homeostasis in ventricular myocytes isolated from mtCaMKII hearts. In order to test if the elevated [Ca^2+^]_i_ in mtCaMKII cardiomyocytes was a consequence of deficient ATP, we fortified the micropipette dialysate with ATP (5 mM) to approximate physiological cytoplasmic ATP activity ^50^. ATP dialysis reduced diastolic [Ca^2+^]_i_ in mtCaMKII cardiomyocytes to WT levels, but had no effect on diastolic [Ca^2+^]_i_ in WT control cardiomyocytes (Fig 3E and F). We interpreted the data up to this point as supporting a model where impaired energetics in mtCaMKII hearts caused defective intracellular [Ca^2+^] homeostasis, a cardinal feature of myocardial dysfunction.

### Replacement of CKmito prevents dilated cardiomyopathy in mtCaMKII mice

Loss of creatine kinase activity is a common finding in failing human myocardium and in animal models of heart failure ^52, 53^. Creatine kinase maintains energetics by rapidly and reversibly producing creatine phosphate, a highly diffusible high-energy phosphate molecule important for cellular energy homeostasis. Over-expression of CK-M, the cytosolic myofibrillar isoform of creatine kinase, in several models of heart failure improves overall cardiac function and reduces cardiac dilation, ^54, 55^ but the potential role of CKmito, the mitochondrial isoform, was only recently shown to improve myocardial response to injury ^56^. We considered that loss of CKmito could be important to the dilated cardiomyopathy in mtCaMKII mice, as it associates with the ATP transporter on the mitochondrial intermembrane space, and enhances energy transfer from the mitochondria to the cytosol ^57^. Furthermore, CKmito was hyperphosphorylated in mtCaMKII compared to WT mitochondria (KCRS, Table S1). We measured expression of myofibrillar, and mitochondrial creatine kinase isoforms in WT and mtCaMKII hearts with antibodies we validated to be specific for each isoform (Fig S5A), and found a trend (P=0.06) for reduced CKmito in mtCaMKII compared to WT (Fig 4A and B). In contrast, the expression of CK-M was similar in WT and mtCaMKII hearts (Fig 4C and D). However, we did not measure a reduction in CKmito 1 week after MI surgery compared to sham operated WT mice (Fig S5B), suggesting mitochondrial CaMKII over-expression in mtCaMKII hearts is a more potent upstream signal for reducing CKmito than MI under these conditions. In order to test whether ATP loss and reduced CKmito contributed to the dilated phenotype in the mtCaMKII hearts, we attempted to recover myocardial energetics by interbreeding mtCaMKII mice with mice over-expressing CKmito in myocardium. CKmito was significantly over-expressed at a similar level in both the CKmito and mtCaMKII x CKmito interbred mice (Fig 4E and F). The mtCaMKII x CKmito interbred mice showed improved in vivo ATP and creatine phosphate levels (Fig 4G), reduced diastolic [Ca^2+^]_i_ in isolated myocytes (Fig 4H), and exhibited repair of dilated cardiomyopathy compared to mtCaMKII littermates (Fig 4I and Fig S2 for cine). Importantly, the recovery of myocardial function and energetics in the mtCaMKII x CKmito mice occurred without a reduction in mitochondrial CaMKII over-expression (Fig 4J and K). In contrast, interbreeding mtCaMKII mice with transgenic mice over expressing the cytosolic CK-M did not rescue dilated cardiomyopathy (Fig S5C), but rather worsened adverse structural remodeling compared to mtCaMKII hearts. These data show that defective energetics and dilated cardiomyopathy in the mtCaMKII mice can be significantly reversed by CKmito replacement. These findings provide new evidence that myocardial dilation can be a metabolic consequence of excessive mitochondrial CaMKII activity leading to ATP deficiency, and suggest that chronic ATP deficiency can result in dilated cardiomyopathy that is potentially preventable or reversible.

**Figure 4.**
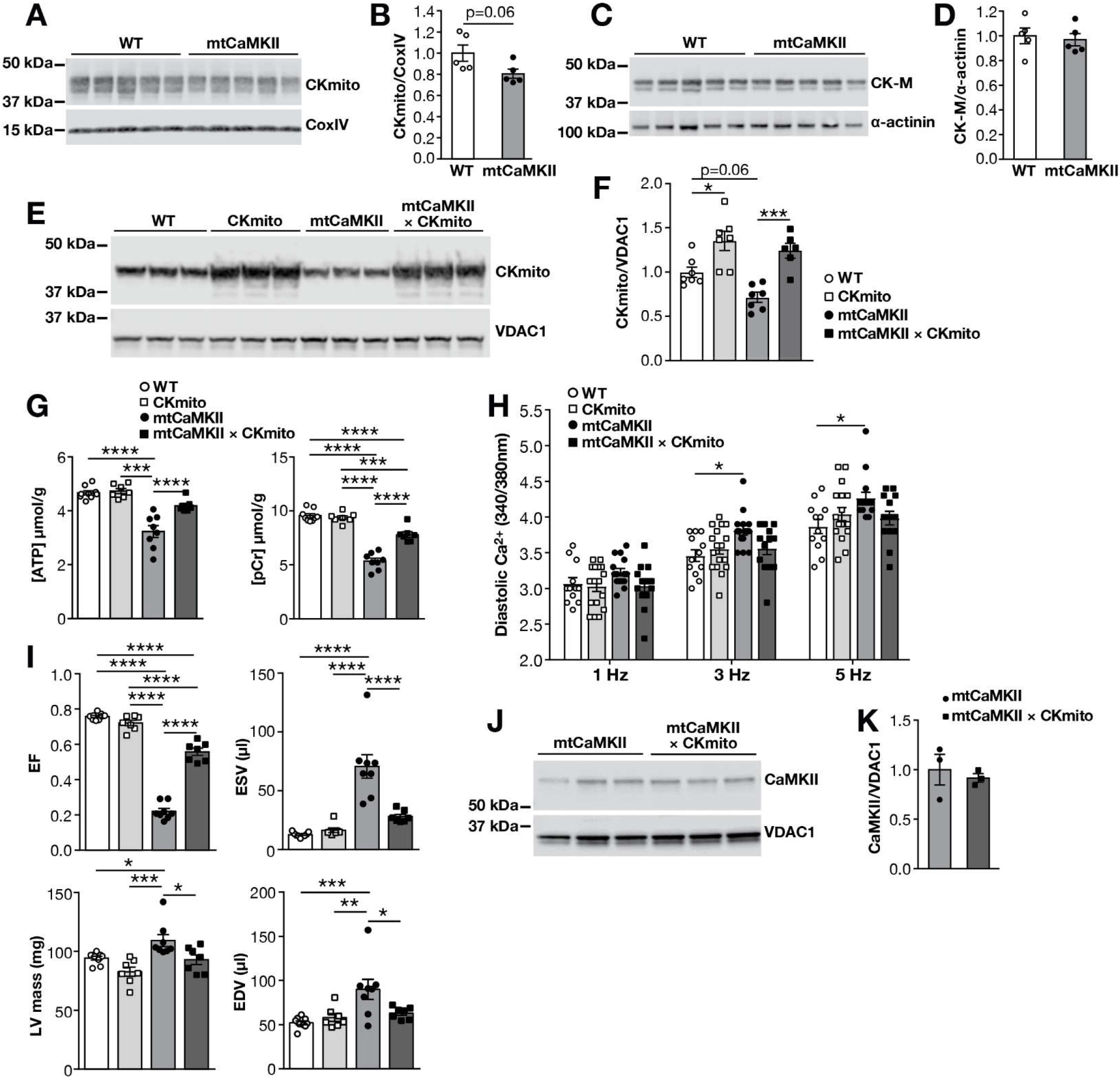
Mitochondrial creatine kinase replacement rescues ATP deficiency in mtCaMKII hearts. (A) Western blots and (B) summary data for CKmito expression normalized to CoxIV from WT (n=5) and mtCaMKII (n=5) hearts. (C) Western blots and (D) summary data for CK-M expression normalized to α-actinin from WT (n=5) and mtCaMKII (n=5) hearts. (E)Western blot and (F) summary data for CKmito expression normalized to VDAC1 from WT (n=7), CKmito (n=7), mtCaMKII (n=7) and mtCaMKII x CKmito (n=6) hearts. (G) In vivo ATP (left) and PCr (right) quantification from ^31^P MR spectroscopy. (H) Summary data for diastolic [Ca^2+^] measurements made with Fura-2–loaded ventricular myocytes stimulated at 1, 3 and 5 Hz (n=12-17 ventricular myocytes isolated from 2 hearts/group). (I) Summary data from in vivo MRI measurements in WT (n=9), CKmito (n=7), mtCaMKII (n=8), and mtCaMKII x CKmito (n=7) hearts. (G-I) WT and mtCaMKII data are the same as in Fig 3. (J) Western blot and (K) summary data for CaMKII normalized to VDAC1 in mitochondrial lysates from mtCaMKII (n=3) and mtCaMKII x CKmito (n=3) hearts. (Data are represented as mean ± SEM, significance was determined using a two-tailed t test or 1 way AVOVA with Tukey’s multiple comparison’s test, ****P<0.0001, ***P<0.001, **P<0.01, *P<0.05)

### Complex I expression and activity are reduced in mtCaMKII hearts

Loss of electron transport chain (ETC) complex components are thought to drive reduced metabolic capacity in some genetic and acquired forms of heart failure ^58, 59^. With this in mind, we next evaluated the ETC to determine if reduced ATP production in mtCaMKII hearts was associated with loss of ETC complex expression and/or function. We interrogated ETC complex expression in heart lysates, and found a significant decrease in protein expression of a complex I component (NDUFB8), and a significant increase in protein expression of a complex II component (SDHB) in mtCaMKII compared to WT hearts (Fig 5A and B). Because these complexes are made up of many proteins, we quantified assembled complexes and measured complex activity using native gel electrophoresis. We found a significant decrease in assembled complex I (Fig 5C) and in complex I activity (Fig 5D) in mtCaMKII mitochondria compared to WT. Based on the apparent decrease in complex I and increase in complex II in mtCaMKII hearts, we measured oxygen consumption rates of isolated mitochondria under conditions selective for complex I (pyruvate/malate) and complex II (succinate/rotenone) metabolism. We observed a decrease in state 3 (ADP activated) respiration in the presence of complex I substrates (pyruvate/malate) (Fig 5E), and an increase in state 3 respiration in the presence of complex II substrates (succinate/rotenone) (Fig 5F) in the mtCaMKII mitochondria compared to WT, consistent with protein expression and activity data. We next repeated the measurement of complex I expression in mtCaMKII mitochondria rescued by CKmito over-expression. We found a partial recovery of expression of a complex I component in mtCaMKII x CKmito compared to mtCaMKII mitochondria (Fig 5G and H), but no recovery of complex I activity (Fig 5I). We interpret these data to collectively indicate that the reduction in ATP production in mtCaMKII hearts is a consequence of extensive metabolic remodeling that can be substantially corrected by a modest replacement of CKmito.

**Figure 5.**
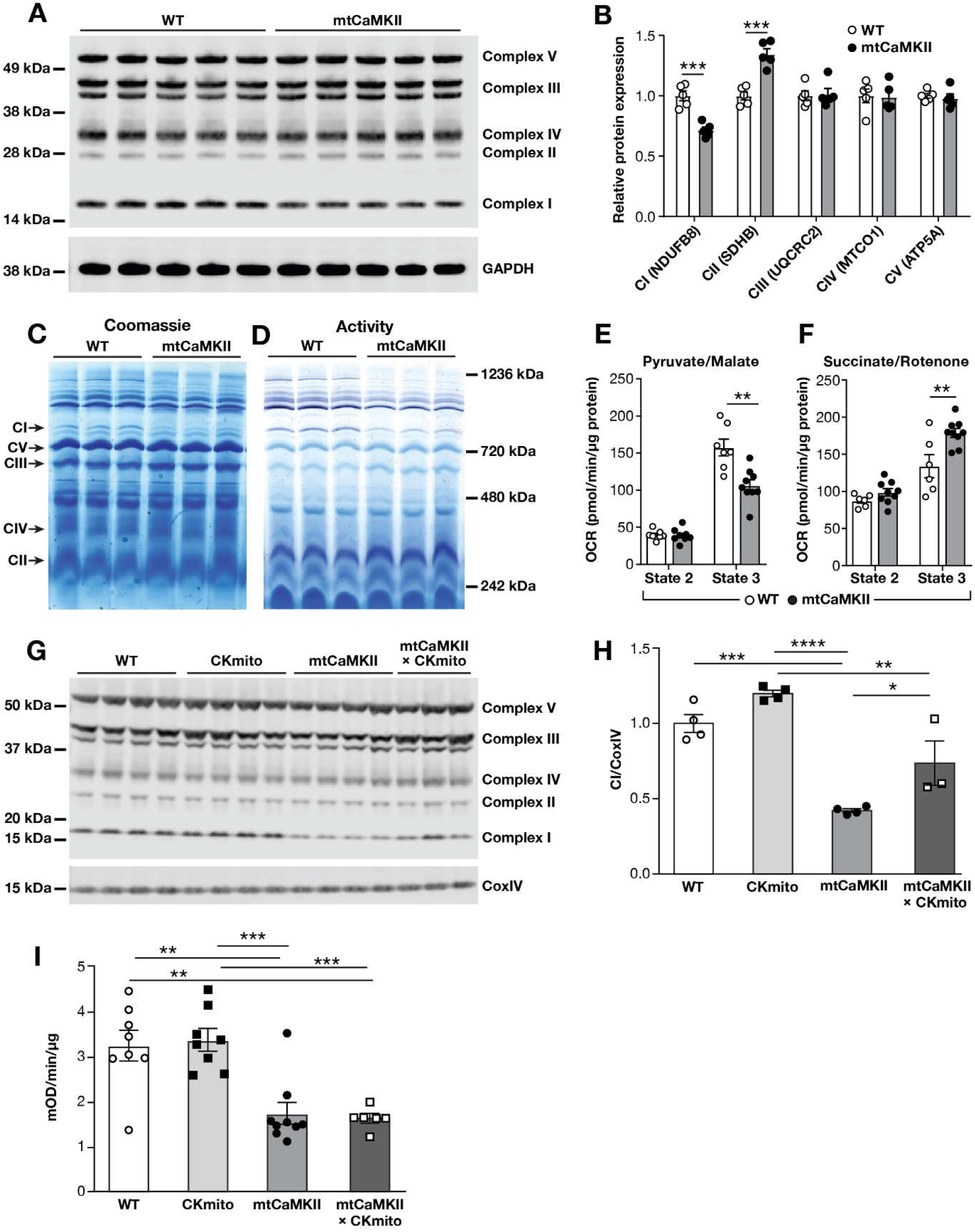
mtCaMKII hearts show decreased complex I and increased complex II activity and expression. (A) Western blot of heart lysates from WT (n=5) and mtCaMKII (n=5) using OxPhos antibody (B) protein expression for complexes I-V normalized to GAPDH. (C) Blue Native gel with mitochondria from WT and mtCaMKII hearts stained with Coomassie blue and (D) in-gel activity assays for complex I. (E and F) Mitochondrial respiration measured under state 2 (substrate alone) and state 3 (substrate plus ADP) conditions in isolated mitochondria from WT (n=7) and mtCaMKII (n=9) hearts. Substrates favored complex I (pyruvate/malate, E) or complex II (succinate/rotenone, F) activity. (G) Western blot with OxPhos antibody and (H) summary data for complex I normalized to CoxIV in heart lysates from WT (n=4), CKmito (n=4), mtCaMKII (n=4) and mtCaMKII x CKmito (n=3) hearts. (I) Complex I activity measured in mitochondria isolated from WT (n=8), CKmito (n=8), mtCaMKII (n=9) and mtCaMKII x CKmito (n=6) hearts. (Data are represented as mean ± SEM, significance was determined using a two-tailed t test or 1 way AVOVA with Tukey’s multiple comparison’s test, ***P<0.001, **P<0.01, *P<0.05)

### Coherent enhancement of TCA cycle dehydrogenases in mtCaMKII hearts

To more broadly interrogate potential metabolic targets for CaMKII in mitochondria we performed a mass spectrometry based analysis of phosphorylated peptides enriched from myocardial mitochondria from mtCaMKII and WT littermate hearts (see Methods). We identified 38 proteins exhibiting significantly increased phosphorylation in the mtCaMKII mice compared to the WT controls. Most notably, 22 of these proteins were involved in metabolism and energy production (Table S1). The abundance of these targets, together with our finding that mtCaMKII have impaired energetics, added further support to our earlier results showing that CaMKII is involved in regulating mitochondrial metabolism.

We were intrigued by a pattern of increased phosphorylation of pyruvate dehydrogenase, and key tricarboxylic acid (TCA) cycle dehydrogenases in the mtCaMKII hearts (Table S1). We next asked if this hyperphosphorylation was associated with a change in activity of these enzymes. The activity of pyruvate dehydrogenase (Fig 6A), α-ketogluterate dehydrogenase, fumarase, and malate dehydrogenase (Fig 6B) were significantly increased in mtCaMKII compared to WT mitochondria, and there was a trend toward increased activity in all TCA cycle enzymes in mtCaMKII compared to WT mitochondria. These findings suggested that mtCaMKII operated, at least in part, to augment metabolism through TCA cycle activity, possibly providing insight into acute metabolic benefits of physiological increases in mitochondrial CaMKII activity. Because the TCA cycle operates to produce NADH, the major electron source for ATP production by oxidative phosphorylation, we measured NADH in ventricular myocytes isolated from mtCaMKII and WT hearts (Fig 6C). Consistent with augmented TCA cycle activity, resting NADH was increased in mtCaMKII compared to WT cardiomyocytes (Fig 6D). However, pacing induced NADH increases were only present in cardiomyocytes isolated from WT mice. In contrast, increased pacing frequency caused a reduction in NADH in mtCaMKII cardiomyocytes (Fig 6D), suggesting that the resting production of NADH was augmented, but lacked capacity to increase NADH consumption during rapid stimulation. Intriguingly, and in contrast to other models of cardiomyopathy with loss of complex I, we did not detect a significant deficiency of NAD+ in cardiac lysates (Fig 6E). NAD+ is an essential cofactor for sirtuins, mitochondrial deacetylases whose loss of function is linked to cardiomyopathy ^60, 61^. We measured an increase in mitochondrial protein acetylation (Fig 6F and G), potentially consistent with previous reports in cardiomyopathy induced by loss of complex I ^62^. However, this enhancement of acetylation was apparently inadequate to cause loss of ΔΨ_mito_ (Fig 2O), or increase cardiomyocyte death (Fig 2M and N), downstream events in cardiomyopathy attributed to NAD+ deficiency and increased mitochondrial protein acetylation ^62^. We interpret these data to indicate excessive mitochondrial CaMKII activity causes profound metabolic remodeling where loss of CKmito is a decisive event leading to myocardial starvation, despite augmented PDH and TCA cycle activity.

**Figure 6.**
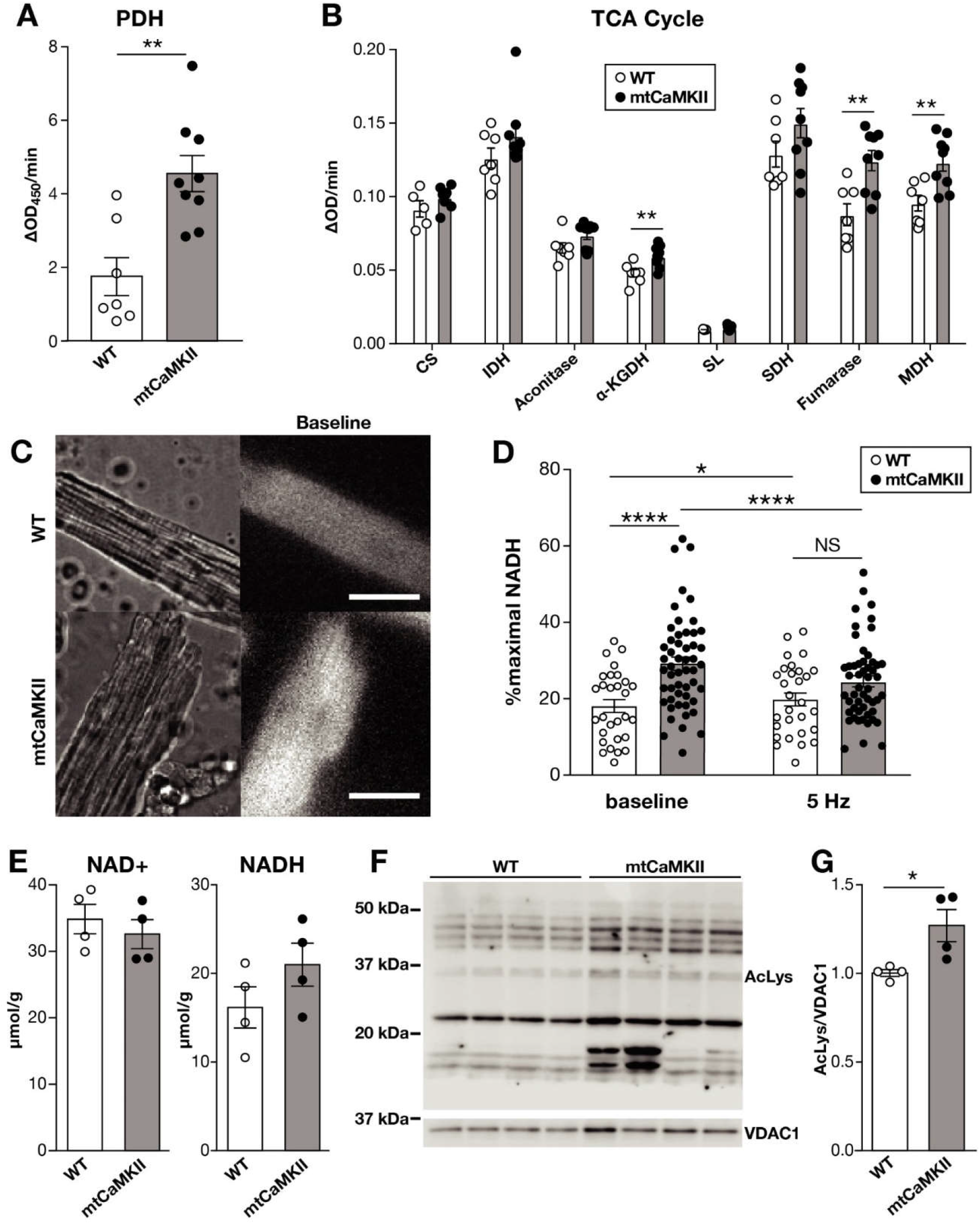
mtCaMKII activates TCA cycle dehydrogenases and augments resting NADH. (A) Pyruvate dehydrogenase (PDH) activity in isolated mitochondria from WT (n=7) and mtCaMKII (n=9) hearts. (B) TCA cycle enzyme activities in permeabilized mitochondria from WT (n=7) and mtCaMKII (n=9) hearts, CS: citrate synthase, IDH: isocitrate dehydrogenase, α-KGDH: α-ketoglutarate dehydrogenase, SL: succinate-CoA ligase, SDH: succinate dehydrogenase, MDH: malate dehydrogenase. (C) Representative NADH imaging of isolated ventricular myocytes before pacing (calibration bar = 50 μm). (D) Quantification of NADH autofluorescence in isolated ventricular myocytes before and after pacing from WT (n=29 myocytes from 2 hearts) and mtCaMKII (n=52 myocytes from 3 hearts) hearts. Maximum (100%) NADH was measured in the presence of NaCN, and minimum (0%) in the presence of FCCP. (E) Quantification of NAD+ and NADH in hearts from WT (n=4) and mtCaMKII (n=4) mice (F) Western blot and (G) summary data for acetylated proteins normalized to VDAC1 in isolated mitochondria from WT (n=4) and mtCaMKII (n=4) hearts. (Data are represented as mean ± SEM, significance was determined using a two-tailed t test, ***P<0.001, **P<0.01, *p<0.05

### Computational modeling predicts loss of ATP, increased NADH and preservation of ΔΨ_mito_ in mtCaMKII hearts

We performed analysis of mitochondrial metabolism using a modified computational model (see Methods) ^63, 64^ (Fig 7A). The effect of mtCaMKII was simulated by reduced CKmito activity and complex I respiration, and increased TCA enzyme activities, as observed in experimental measurements (Table S2). This mathematical simulation showed decreased ATP and creatine phosphate levels in mtCaMKII, and replacement of CKmito produced a substantial rescue of these parameters (Fig 7B and Fig S6), similar to our experimental observations. We interpret the consistency between these simulated and experimental outcomes to suggest that the combination of reduced CKmito and complex I activity are sufficient to cause decreased ATP, despite enhanced pyruvate dehydrogenase and TCA cycle activity, and that ATP deficiency can be repaired by CKmito replacement in the presence of excessive mtCaMKII.

**Figure 7.**
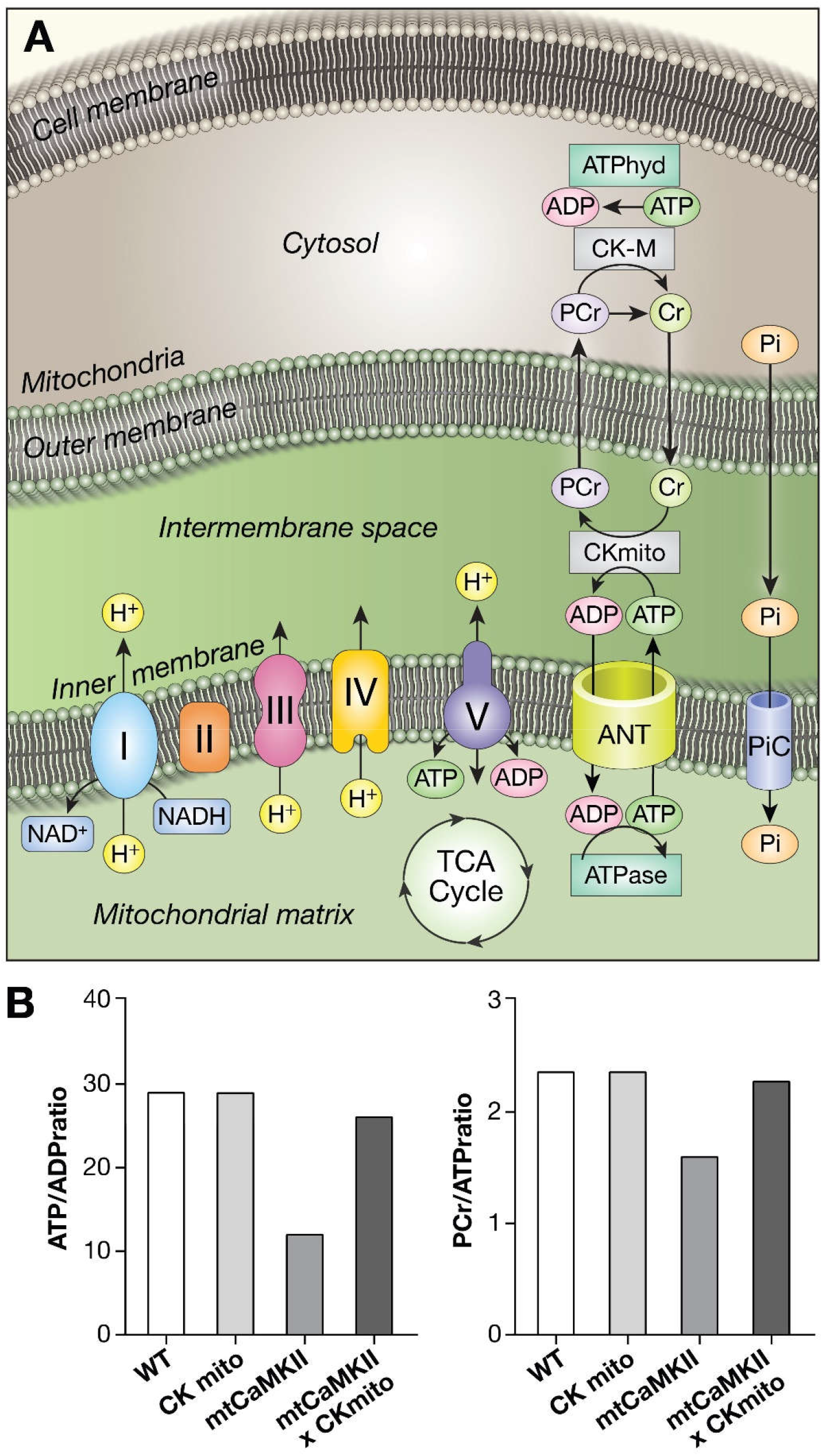
Computational modeling of energetics. (A) Schematic image of key parameters used in a three compartment model of CaMKII, creatine kinase (CK), and mitochondrial energetics. (B) Computer stimulation results were compared in WT, CKmito, mtCaMKII, and mtCaMKII x CKmito interbred conditions. The ratios of PCr/ATP and ATP/ADP were calculated at the end of 5 minutes cardiac cycle (simulated using a pulsatile function to oscillate ATP hydrolysis).

## Discussion

### Failing myocardium and failing energetics: chicken or egg?

The association between defective ATP metabolism and heart failure has been recognized for many years ^65, 66, 67^, and diminished expression of creatine kinase and complex I have been noted in animal models, and in failing human hearts ^53, 54, 68, 69, 70^. Despite this, there has been uncertainty whether failed energetics are a cause or a consequence of damaged myocardium. Thus, lack of understanding of upstream signals controlling energetics and cardiomyopathy constitutes an important knowledge gap. Our study provides new evidence that mitochondrial CaMKII can orchestrate pathological metabolic remodeling and left ventricular dilation. mtCaMKII hearts are deficient in ATP (Fig 3C), dilated and manifest contractile dysfunction and deranged Ca^2+^ homeostasis, but are free of hypertrophy and fibrosis. We acknowledge that the derangement in energetics could be secondary to the severe pathology we observed in the mtCaMKII hearts since we were unable measure ATP in hearts prior to the development of these pathological phenotypes due to technical limitations. However, excessive mitochondrial CaMKII activity is almost certainly the inciting trigger due to the nature of mitochondrial-targeted CaMKII over-expression in this model. In contrast to mice with untargeted myocardial CaMKII over-expression, the mtCaMKII do not show a high rate of premature death, despite depressed energetics and cardiac chamber dilation. Restoration of a single enzyme, CKmito, by interbreeding mtCaMKII with CKmito transgenic mice, substantially repaired energetics and cardiac dysfunction, and restored [Ca^2+^]_i_ homeostasis (Fig 4G-I). Additionally, [Ca^2+^]_i_ homeostasis can be restored by supplementing ATP directly to the cytosol of myocytes (Fig 3E and F). These findings are corroborated by our computational modeling where replacing CKmito activity improved ATP levels in the mtCaMKII model (Fig 7 and Fig S6). These data suggest that the lack of adequate energy can directly lead to a stable pattern of ventricular dilation and contractile dysfunction, and restoring myocardial energy supply significantly improves cardiac function, despite persistence of excessive mitochondrial CaMKII activity.

Our computational modeling confirmed that ATP deficiency in mtCaMKII hearts is also partly a consequence of loss of complex I activity, and persists despite hyperactivation of the TCA cycle, and increased NADH. Our phosphoproteomic studies identified a novel site for PDH phosphorylation that is different from the well-known inhibitory sites regulated by mitochondrial calcium ^71^. We observed enhanced PDH activity in the mtCaMKII hearts, but further investigation will be required to know if this novel phosphorylation site contributes to PDH activation under conditions of excessive CaMKII activity. Diminished complex I content is noted in ischemic and genetic forms of cardiomyopathy ^68, 70^. Our phosphoproteomic studies also identified NDUFB11, a complex I subunit essential for complex assembly and stability ^72, 73^, as a CaMKII phosphorylation target, raising the possibility that excessive mitochondrial CaMKII activity has adverse consequences for complex I assembly. Other models of complex I deficiency show reduced sirtuin activity due to loss of the cofactor, NAD+, and it is possible that increased complex II activity was protective against excessive loss of NAD+ in mtCaMKII hearts compared to other models marked by loss of complex I. Loss of sirtuin mediated deacetylase activity results in excessive acetylation of mitochondrial proteins, mPTP opening, increased ROS, and myocyte death ^62^. We did not find a significant deficiency of NAD+ in mtCaMKII hearts, and the increases in acetylation in the mtCaMKII mitochondria were evidently insufficient to trigger loss of ΔΨ_mito_ and cell death. Further studies will be needed to understand how reduced ATP, elevated [Ca^2+^]_i_, and, perhaps, other factors related to excessive mitochondrial CaMKII activity lead to left ventricular dilation. It does seem clear that the increased diastolic [Ca^2+^]_i_ observed in the mtCaMKII cardiomyocytes was insufficient to substantially activate Ca^2+^ sensitive hypertrophic signaling pathways. Our current findings suggest mitochondrial CaMKII can contribute to pathological injury by remodeling metabolism.

### CaMKII directs cardiomyopathy phenotypes by subcellular domain specific actions

Our results support a view that organelle resident and subcellular specific actions of CaMKII can selectively drive distinct cardiomyopathy phenotypes. Nuclear CaMKII over-expression causes mild myocardial hypertrophy ^12^, and non-targeted CaMKII over-expression leads to massive hypertrophy, myocardial death, fibrosis, chamber dilation, heart failure, deranged Ca^2+^ homeostasis, and early sudden death ^13, 51^. Here we show that targeted mitochondrial CaMKII over-expression causes a surprisingly pure dilated cardiomyopathy linked to adverse metabolic remodeling, and ATP deficiency. Unexpectedly, chronic actions of excessive mtCaMKII do not cause increased myocardial or sudden death. We have not yet identified a mechanism for CaMKII translocation to mitochondria, but note, similar to the case of CaMKII, that multiple mitochondrial proteins lack canonical TOM/TIM guide sequences ^74^. We assume mitochondrial CaMKII is activated by Ca^2+^/calmodulin, ROS and O-GlcNAcylation, upstream CaMKII activating signals that are present in mitochondria ^75, 76^. The sequence of compartment specific increases in CaMKII activity in heart following injury are unknown, but our findings suggest that cellular stress pathways, activated by MI, likely contribute to increased mitochondrial CaMKII activity. The potential for CaMKII to operate in various cellular compartments suggests that CaMKII functions as a highly versatile agent, promoting hypertrophy and cardiac chamber dilation, depending on its subcellular location.

### Failed myocardial energetics impair diastolic Ca^2+^ homeostasis in mtCaMKII mice

Failing human myocardium shows elevated diastolic [Ca^2+^]_i_ and increased cytosolic CaMKII activity, as evidenced by ryanodine receptor hyperphosphorylation and sarcoplasmic reticulum Ca^2+^ leak ^77^. Acute CaMKII inhibition is sufficient to reduce Ca^2+^ leak, lower diastolic [Ca^2+^]_i_, and improve mechanical function in failing human myocardial strips ^77^. Our studies extend the connections between CaMKII and cellular Ca^2+^ homeostasis by demonstrating that loss of cytoplasmic [Ca^2+^]_i_ homeostasis and left ventricular dilation can arise from defective myocardial energetics, independent of increased cytoplasmic CaMKII activity because cardiac chamber dilation persisted in mtCaMKII mice interbred with AC3-I transgenic mice with extramitochondrial CaMKII inhibition (Fig 2C). Myocardium is highly dependent on a moment to moment balance of ATP production and utilization related, in part, to the high energy demands of continuous and physiologically changing work ^78^. Achieving physiological [Ca^2+^]_i_ homeostasis is a major energetic task for myocardium ^79^, and elevation of diastolic cytoplasmic [Ca^2+^] is a common finding in animal heart failure models ^80^ and in failing human myocardium ^81^, conditions where energy supply is insufficient to satisfy demand. The mtCaMKII hearts show a severe loss of ATP and reduction in CKmito and complex I, consistent with a scenario where energy stores are inadequate to support normal function. ATP flux is purposed to specific subcellular compartments, and mitochondrial ATP is primarily responsible for fueling SERCA2a activity that is required to stabilize diastolic cytoplasmic [Ca^2+^] after each heartbeat ^82, 83^. Myofilament activation and cardiac contraction follow a regenerative release of Ca^2+^ to the cytoplasm from the intracellular sarcoplasmic reticulum store, while relaxation and diastole are a consequence of sequestration of cytoplasmic Ca^2+^ into the sarcoplasmic reticulum by SERCA2a. We found that decreased ATP in mtCaMKII hearts was associated with elevation of diastolic cytoplasmic [Ca^2+^], and that elevated [Ca^2+^] could be repaired in vitro by exogenous ATP and in vivo by replacement of CKmito. We interpret these findings to provide strong support for the hypothesis that mtCaMKII dilated cardiomyopathy arises from ATP deficiency that is exacerbated by CKmito deficiency. We speculate that sustained hyperactivation of mitochondrial CaMKII could contribute to metabolic defects and elevated diastolic [Ca^2+^] in human heart failure. Additionally, others have shown that increases in myocardial ADP and intracellular [Ca^2+^] can contribute to diastolic dysfunction^84^. While we did not directly quantify diastolic function, this may also be contributing to the phenotype of the mtCaMKII mice.

### What is the relationship of LV dilation to LV hypertrophy in pathological myocardial remodeling?

The core components of adverse cardiac remodeling are myocardial hypertrophy and dilation ^26, 27^. However, the biological connections between hypertrophy and dilation are uncertain. One paradigm is that myocardial hypertrophy is a compensatory reaction to pathological stress, but sustained and excessive hypertrophy can progress to chamber dilation over time. Although the interrelationship between hypertrophy and dilation is not completely understood, dilation may be favored by myocyte death interposed upon hypertrophied myocardium that eventually impairs myocardial ability to sustain elevated ventricular wall stress. However, the rate of progression to left ventricular dilation in patients with left ventricular hypertrophy detectable by sonographic imaging is low ^85^, suggesting that the processes of hypertrophy and dilation are not tightly coupled, nor necessarily driven by the same biological mechanisms. Our findings regarding mtCaMKII cardiomyopathy indicate that myocardial dilation can arise as a sustained response to myocardial energy deprivation in the absence of myocardial hypertrophy, or death. The cardiac dilation in mtCaMKII hearts is potentially reversible, because it does not involve increases in myocardial death, or fibrosis, and can be rescued by replacement of CKmito. Furthermore, [Ca^2+^]_i_ homeostasis can be repaired in ventricular myocytes isolated from mtCaMKII hearts by addition of ATP. Thus, it is possible that CKmito replacement, or mitochondrial CaMKII inhibition could prevent or reverse some acquired forms of dilated cardiomyopathy, a life-threatening disease phenotype that has been largely considered irreversible.

### What is the physiological role of mitochondrial CaMKII?

Many studies have provided evidence affirming a role of excessive CaMKII in myocardial disease. However, relatively less is known about the physiological roles of CaMKII in heart. Surprisingly, CaMKII activity appears dispensable for basal cardiac function ^15, 22^, although complete ablation of CaMKIIγ and CaMKIIδ in myocardium causes mild hypertrophy due to loss of a tonic negative regulatory action on calcineurin signaling ^86^. We interpret the available evidence to suggest that CaMKII promotes fight or flight responses that increase myocardial performance during extreme physiological stress. Many of these responses are linked to CaMKII catalyzed phosphorylation of Ca^2+^ homeostatic proteins that enhance the excitation-contraction coupling mechanism. For example, CaMKII may augment the treppe effect in ventricular myocardium ^87^, and enhance the dynamic range of heart rate acceleration in cardiac pacemaker cells ^88^ by catalyzing ryanodine receptor phosphorylation. Our new findings showing enhanced TCA cycle activity indicate that mitochondrial CaMKII can increase delivery of NADH to the electron transport chain. While we acknowledge that chronic over-expression of CaMKII creates an artificial condition, and although we currently lack tools to tightly control mitochondrial CaMKII expression on a moment to moment time scale, we speculate that acute, physiological activation of mitochondrial CaMKII could be advantageous by increasing ATP flux from oxidative phosphorylation by enhanced NADH production through the TCA cycle. However, mtCaMKII mice, burdened by chronic, pathological elevation in mitochondrial CaMKII, exhibit loss of CKmito and complex I that prevent this metabolic benefit, and instead contribute to metabolic insufficiency, elevated diastolic cytoplasmic [Ca^2+^], and severe dilated cardiomyopathy.

## Methods

### Animal Models

All the experiments were carried out in accordance with the guidelines of Institutional Animal Care and Use Committee (PHS Animal Welfare Assurance, A3021-01 (Univ. Iowa), A3272-01 (JHU)). Mice used in these studies were a mixture of male and female animals 7-20 weeks of age unless otherwise noted in the figure legend.

### Myocardial Infarction

Mice are anesthetized with 1.5-2% isoflurane and given a pre-emptive dose of buprenorphine (0.03-0.07mg/kg). Mice are then intubated and ventilated (100% O_2_, 200ul TV, 120 bpm) using a Harvard Apparatus MiniVent. When a toe pinch confirms proper depth of anesthesia, mice are given a single dose of succinylcholine (2mg/kg). Body temperature is maintained at 37°C using a rectal thermometer and infrared heating lamp. A left thoracotomy and pericardiotomy is performed, and the left main coronary artery is completely occluded by tying a knot with 7-0 prolene. After verification that MI has occurred (blanching of the tissue distal to the suture), the ribs and skin are closed with 5-0 silk. Once the chest is closed, the mouse remains under anesthesia until succinylcholine is completely metabolized. Isoflurane is turned off, the mouse is allowed to regain respiration on its own, and then extubated. When performing a sham surgery, mice undergo the same procedure, but the ligation of the artery is excluded. A second dose of buprenorphine (0.06-0.075mg/kg) is given before returning mice to animal facility. The surgeon was blinded to the genetic identity of the mice for all studies.

### TTC staining and MI sizing

At the end of 24 hours post-MI, mice were sacrificed and the heart was quickly excised, and both atria and right ventricular free wall were removed. Left ventricular tissue was weighed and wrapped with cling film and then frozen at −20°C for 45 min, and then sliced into 6-7 pieces of 1.0 mm thick sections perpendicular to the long axis of the heart. The sections were incubated individually using a 24-well culture plate in 1% TTC in phosphate-buffered saline at pH 7.4 at 37°C for 10 min), and then digitally photographed. For infarct size at 24 hours post-MI, TTC-stained area, and TTC-negative staining area (infarct myocardium) were measured using ImageJ. Myocardial infarct size was expressed as a percentage of the total LV area.

### mtCaMKII mice

The mouse CaMKIIδ_C_ cDNA was fused with an N-terminal flag epitope tag and the N-terminal 28 amino acids of cox8a for mitochondrial targeting. The resulting construct was cloned into the pBS-αMHC-script-hGH vector for myocardial expression. Pronuclear injections of linearized DNA (digested with NotI) were performed in the University of Iowa Transgenic Mouse Core Facility and embryos implanted into pseudo-pregnant females to generate C57Bl6/J F1 mice. Insertion of the transgene into the mouse genome was confirmed by PCR analysis (not shown) using the forward primer, 5’- GCA GTC AGA AGA GAC GCG -3’, and reverse primer, 5’- GAA TCC CAA CAA CTC GGG AGG C -3’, producing a product of 500 bases.

### CKmito mice

The CKmito transgenic mice were generated using aHMC-Tet promoter ^89^ fused with CKmito cDNA and human growth hormone polyA. The resulting TetCKmito mice were bred with aMHC-Tet-off mice to generate compound heterozygous mice as the CKmito transgenic mice ^89, 90^. Mouse genotypes were determined using real time PCR with specific probes by a commercial vendor (Transnetyx, Cordova, TN). The CKmito transgenic mice express CKmito specifically in cardiomyocytes in the absence of doxycycline treatment.

### Echocardiography

Transthoracic echocardiography was performed in unsedated mice using a Vevo 2100 (VisualSonics Inc) system, equipped with a 30-40 MHz linear array transducer. The left ventricle end-diastolic and end-systolic ventricular volumes (EDV, ESV), and the percent ejection fraction (EF) were estimated using the Simpson’s method from the apical two chamber view of the heart ^91, 92^. Measurement of EDV, ESV were manually traced according the American Society of Echocardiography ^93^. The EF was automatically calculated by the system building software using the following formula:

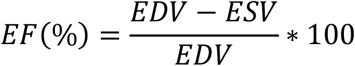

The M-mode echocardiogram was acquired from the parasternal short axes view of the left ventricle (LV) at the mid-papillary muscles level and at sweep speed of 200 mm/sec. From this view left ventricle mass (LV mass) is calculated from inter-ventricular septal thickness at end of diastole (IVSD), LV chamber diameter at end of diastole (LVEDD), posterior wall thickness at end of diastole (PWTED) measurements using the following formula where 1.055 is the specific gravity of the myocardium ^94^:

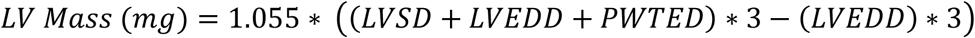

The echocardiographer was blinded to the genetic identity of the mice for all studies.

### In vivo Murine Cardiac MRI/MRS

Anesthesia was induced under 2% isofluorane and maintained under 1% isofluorane and the in vivo studies carried out on a Bruker Biospec MRS/MRI spectrometer equipped with a 4.7T/40cm Oxford magnet and a 12cm (i.d.) actively shielded BGA-12™ gradient set, as previously described ^50, 54^. <u>MRI for Ventricular Size and Function</u>: A complete set of high temporal and spatial resolution multi-slice cine images were acquired and analyzed as previously reported ^49, 95, 96^. LV EF was calculated from the difference in end-diastolic and end-systolic cavity volumes divided by the end-diastolic volume. <u>^31^P MRS protocol</u>: After shimming, a one-dimensional ^31^P chemical shift imaging (1D-CSI) sequence, using adiabatic excitation pulses to obtain uniform flip angle across the field of view, was acquired as previously described ^50^. The PCr and [β-P]ATP peaks in ^31^P MR localized spectra are quantified by integrating peak areas ^49, 95, 96^. Voxel shifting, a processing method to shift the slice boundaries fractionally, permits the redefiniton of the slice boundaries after data collection to align them with the LV border and minimize contamination from the chest wall, using high resolution ^1^H images as a guide ^97, 98^. Concentrations of PCr and ATP as well as the ratio of PCr/ ATP were measured as previously described ^50^. The MRI studies were performed without knowledge of the genetic identities of the mice.

### Western blot

Heart lysates were prepared in RIPA buffer containing protease and phosphatase inhibitors. Heart lysate or cytoplasm and mitochondria fractions were loaded into 4-12% Bis-Tris NuPAGE gels from (Thermofisher) or 4-15% Mini Protean gels (BioRad). Proteins were transferred to nitrocellulose membrane using Turbo-blotter (Bio-Rad). Primary antibodies [CaMKII (Abcam), phospho-CaMKII (Thermo Scientific), VDAC1(Abcam), FLAG (Rockland), GAPDH (Cell Signaling), CKmito (Sigma), CoxIV (Cell Signaling), CK-M (Sigma), α-actinin (Sigma), OxPhos (Abcam), AcLys (Abcam)] were incubated with the membrane overnight at 4°C. Secondary antibodies were incubated with the membrane at room temperature for 1 hour. Blots were imaged using an Odyssey Fc Imager (Licor). Bands were quantified using Image Studio Software (Licor).

### Mitochondrial isolation

Mitochondrial were isolated using homogenization method on ice as previously described ^99^. In brief, the heart was minced and homogenized in ice-cold isolation buffer (75 mM sucrose, 225 mM mannitol, 1 mM EGTA, and 0.2% fatty acid free BSA, pH 7.4) using a Potter-Elvehjem glass homogenizer. The homogenates were centrifuged for 10 min at 500Xg. The resulting supernatants were centrifuged for 10 min at 10,000Xg. The mitochondrial pellets were washed twice in isolation buffer and centrifuged at 7,700Xg for 6 min each. Mitochondrial pellets were suspended in a small amount of isolation buffer to ~5mg/ml. The protein concentration determined by using the BCA Assay kit (Thermo Fisher Scientific).

### Mitochondrial ATP measurements

Mitochondrial ATP content was measured using ATP Bioluminescence Assay Kit CLS II, Roche Life Science). Isolated mitochondria (15 μg) were suspended in KCl based buffer at 37°C. The ATP production was calculated from a standard curve for a series of ATP concentrations.

### Mitochondrial Ca^2+^ content

Total mitochondrial Ca^2+^ content was measured in isolated mitochondria with the Calcium Assay Kit (Cayman Chemical). Total Ca^2+^ was normalized to protein concentration.

### Mitochondrial ROS production

Amplex UltraRed, a H_2_O_2_ sensitive fluorescent probe, was used to monitor ROS production in isolated mitochondria. Hydrogen peroxide was used to establish the standard curve as previously described ^100^. Briefly, 15 μg of isolated mouse heart mitochondria was suspended in 0.2ml KCl based buffer solution (137 mM KCl, 0.25 mM EGTA, 2 mM MgCl_2_, 2mM KH_2_PO_4_, 20 mM HEPES, 5 mM NaCl, pH 7.2) with 4 U/ml horseradish peroxidase, 40 U/ml superoxide dismutase, 10 μM Amplex UltraRed in a 96-well plate, followed by sequential additions of substrate 5mM pyruvate/malate, complex I inhibitor 0.5 μM rotenone and complex III inhibitor 1 μg/ml anitimycin A. Amplex UltraRed fluorescent signal was read by a microplate reader at 535nm excitation and 595nm emission.

### Mitochondrial membrane potential

Mitochondrial membrane potential (ΔΨ_mito_) was monitored by the ratiometric dye tetramethylrhodamine methyl ester (TMRM) (546/573nm excitation and 590nm emission). Isolated mitochondria (25 μg) were suspended in 0.2ml KCl based buffer solution (137 mM KCl, 0.25 mM EGTA, 2 mM MgCl_2_, 2mM KH_2_PO_4_, 20 mM HEPES, 5 mM NaCl, pH 7.2) in a 96-well plate and read by using a BioTek Microplate reader. Glutamate/malate (5mM/5mM) were used as substrates to energize mitochondria (state 2) follow by addition of 1mM ADP (state 3).

### Blue Native Gel and Complex I activity

Mitochondrial isolates from heart tissue lysates were prepared according to the NativePAGE Sample prep kit (ThermoFisher) (8g/g: digitonin/protein). The mitochondrial resuspensions were run on 3-12% Bis-Tris Native Page gels (ThermoFisher). Gels were run using the NativePAGE Running Buffer kit (ThermoFisher). Gels were either stained using the Coomassie R-250 Staining kit (ThermoFisher) or used for in-gel activity. For this, the gel was equilibrated in 5 mM Tris HCl, pH 7.4 for 10 min and then developed in complex I activity solution (5 mM Tris pH7.4, 2.5mg/ml NBT, 0.1mg/ml NADH). The reaction was terminated by soaking the gel in 10% acetic acid. Gels were imaged using an Epson Perfection V800 Photo scanner. For the microplate plate assay, Complex I activity was measured using the Complex I Enzyme Activity Microplate Assay Kit (Abcam ab109721) as specified by the manufacturer’s instructions. Briefly, 100 μg mitochondrial pellets were resuspended in 25 μL protein extraction buffer and incubated on ice for 30 minutes. Samples were then centrifuged at 16,000Xg for 20 minutes at 4°C. Supernatant protein concentration was determined via BCA Assay kit (Thermo Fisher Scientific). 20 μg of protein were combined with Incubation Solution (Abcam) to a total of 1 mL. Each sample was loaded in triplicate (200 μL/well) and incubated for 3 hours at room temperature. Complex I activity was determined by following change in 450 nm absorbance per minute (mOD/min) for 30 minutes following the addition of Assay Solution (Abcam).

### Mitochondrial Respiration/Oxygen consumption

Mitochondrial oxygen consumption rate was assayed using a high-throughput automated 96-well extracellular flux analyzer (XF96; Seahorse Bioscience). Freshly isolated mitochondria (1-5μg of mitochondrial protein) were suspended in the potassium-based buffer (137 mM KCl, 2 mM KH_2_PO_4_, 0.5 mM EGTA, 2.5 mM MgCl_2_, and 20 mM HEPES at pH 7.2, and 0.2% fatty acid free BSA), transferred into a 96-well XF96 plate, and centrifuged at 3,000Xg for 20 min at 4°C to attach to the plate. The respiration was evaluated with substrates of complex I (5mM pyruvate/malate), complex II (5mM succinate), following addition of 1 mM ADP. Oxygen consumption was measured with one cycle of 0.5 min mix, 3 min measurement and another 0.5 min mix in each step after port injection ^101^.

### TCA Cycle enzyme activities

Pyruvate dehydrogenase (PDH) activity was measured with the Pyruvate Dehydrogenase Enzyme Activity Microplate Assay Kit (Abcam) using 20μg of isolated mitochondria for each assay. Citrate synthase (CS) activity was measured using a MitoCheck Citrate Synthase Activity Assay Kit (Cayman Chemicals). The rest of TCA cycle enzymes activities were measured using colorimetric absorbance based kinetic assays as described ^102^. Briefly, the isolated mitochondria were freeze/thawed 3 times to lyse the mitochondria. Mitochondrial lysate (15 μg) was then suspended in phosphate based solution (50mM KH_2_PO_4_ with 1mg/ml BSA, pH7.2 in the first assay, 10mM KH_2_PO_4_ in the second assay). The first assay used dichlorophenol Indophenol (DCPIP) reduction (λ_absorbance_=600nm) to measure succinyl-CoA ligase (SL), succinate dehydrogenase (SDH), fumarase (FM) and, malate dehydrogenase (MDH) by sequential addition of reagents 140μM DCPIP, 100 μM Duroquinone, 0.8mM Phenazine Methosulfate (PMS), 8μM rotenone/100 μM succinyl CoA and 02.mM KCN, 10mM succinate, 10mM malonate, 10mM glutamate plus 800 μM NAD+ and 1IU (AAT), 15mM fumarate, and finally 10mM malate. The second spectrophotometric assay measures α-ketoglutarate dehydrogenase (KGDH), aconitase (AC) and isocitrate dehydrogenase (IDH) by monitoring NADH/NADPH reduction in 800μM NAD+, 2mM Dithiothreitol (DTT), 100uM EDTA, 2mM CaCl_2_, 2mM MgCl_2_, 2mM α-ketoglutarate, 100μM thiamine pyrophosphate (TPP) and 0.1% Triton X100 by sequential additions of 0.5mM CoASH, 0.8mM NADP+, 5mM MgCl_2_, 0.5mM cis-aconic acid and 15mM isocitrate. Absorbance measurements were made using a BioTek Microplate reader (Synergy MX 96 well plate reader).

### Adult ventricular myocyte isolation

Adult ventricular myocytes were isolated as previously described ^103^. Briefly, mice (8-12 weeks, either gender) were anesthetized by Avertin injection. Hearts were rapidly excised and placed in ice cold nominally Ca^2+^ free HEPES-buffered Tyrode’s solution. The aorta was cannulated, and the heart was perfused in a retrograde fashion with a nominally Ca^2+^ free perfusate for 5 min at 37°C. This was followed by a 15-min perfusion with collagenase-containing nominally Ca^2+^ free solution. Final perfusion was with collagenase-containing low Ca^2+^ (0.2 mM) solution. The LV and septum were cut away, coarsely minced and placed in a beaker containing low Ca^2+^ solution with 1% (w/v) BSA at 37°C. Myocytes were dispersed by gentle agitation, collected in serial aliquots and then maintained in standard saline solution containing 1.8 mM Ca^2+^.

### Immunofluorescence of isolated caridac myocytes

Isolated ventricular myocytes were fixed in 100% ethanol prior to incubation with antibodies Flag (Rockland), CoxIV (Cell Signaling). Stained cells were mounted on slides, and imaged on a laser-scanning confocal microscope (LSM 510, Carl Zeiss).

### Immunofluorescence of heart sections

In brief, hearts were fixed in 4% PFA and embedded in OCT freezing compound (Fisher Scientific, Waltham MA, USA). 10 μm cardiac two chamber view sections were cut using a Microm HM 550 cryostat. Sections were post-fixed with 4% paraformaldehyde and permeabilized using 0.1% Triton-X in PBS. Sections were washed two times for 5 minutes with 1× HBSS. Sections were incubated with anti-CD45 (Thermo Fisher, Waltham, MA, USA) (#14-0451-82 at 1:50) according to previously published methodology ^104^. Samples were subsequently incubated with Alexafluor 555 at 1:1000 (Thermofisher). At least 3-5 fields of view per infarct and remote zone were assessed from n=5 hearts per genotype. Images were acquired in 0.63 μm Z-stacks using an Olympus IX83 microscope, 60×/1.42NA PLAPON objective. CD45+, nucleated cells were counted and normalized number of cells/area (mm^2^).

### Myocyte cross-sectional area

In brief, hearts were prepared as above. Sections were incubated with 5 μg/mL in Wheat Germ Agglutinin, Alexa Fluor^®^ 488 conjugate (Molecular Probes, Eugene, OR, USA) for 20 minutes. Samples were washed two times for 5 minutes with 1× HBSS and mounted with Prolong Diamond Anti-Fade Reagent (Molecular Probes, Eugene, OR, USA). Ten fields of view were acquired from each animal. At least four animals were processed per genotype. 0.63 μm Z-stacks were collected using an Olympus IX70 microscope, 40×/0.75NA UPLFLN-PH objective, and deconvolved using constrained iterative deconvolution software in the cellSens software suite. Average projections for each field of view were calculated and analyzed using cellProfiler (Cell Profiler Inc. Boston, MA, USA) according to previously published methods to determine cell size.

### Mitochondrial Ca^2+^ uptake

Mitochondrial Ca^2+^ uptake was measured as previously described (Joiner, 2012) in saponin-permeablized ventricular myocytes. Assays were performed with freshly isolated myocytes in a 96-well assay plate in respiration buffer (100 mM KAsp, 20 mM KCl, 10mM HEPES, 5mM glutamate, 5mM malate, and 5mM succinate, pH 7.3) supplemented with 100 μM blebbistatin, 5 μM thapsigargin, 0.005% saponin, 5 μM CsA and 1 μM Ca^2+^ green-5N (CaG5N, Thermo Scientific). CaG5N fluorescence was monitored (485 nm excitation, 535 nm emission) after adding CaCl_2_ (50 μM free Ca^2+^) at 3 min intervals at 30°C. Cells from WT and mtCaMKII hearts were prepared on 3 separate days, and the mean of measurements read from 4 wells reported.

### Cytosolic Ca^2+^ Measurements

Cytosolic Ca^2+^ levels were recorded from Fura-2–loaded ventricular myocytes as previously described ^82^. Briefly, single isolated ventricular myocytes were loaded with 4 μM Fura-2 acetoxymethyl (AM) for 30 min and then perfused with Tyrode’s solution for 30 min to de-esterify the Fura-2 AM. After placement on a recording chamber, the cells were perfused in bath solution comprised (mM): 137 NaCl, 10 Hepes, 10 glucose, 1.8 CaCl_2_, 0.5 MgCl_2_, and 25 CsCl; pH was adjusted to 7.4 with NaOH, at 35 ± 0.5 °C. Whole-cell patch clamp technique was used to dialyze cells. The pipette (intracellular) solution comprised of (mM): 120 CsCl, 10 Hepes, 20 TEA chloride, 1.0 MgCl_2_, 0.05 CaCl_2_, 10 glucose; the pH was adjusted to 7.2 with 1.0 N CsOH. Adenosine 5’-triphosphate disodium salt hydrate 5 mM was added to the pipette solution when needed. Myocytes were stimulated at 1, 3 and 5Hz using a voltage protocol holding at −80 mV and stepping to 0 mV for 100 ms from a prepulse of 50 ms at −50 mV. The cytosolic Ca^2+^ transients were measured from cells excited at wavelengths of 340 and 380 nm and imaged with a 510-nm long-pass filter.

### NADH measurements

The autofluorescence of endogenous NADH was measured as described ^83^. Briefly, ventricular myocytes were put into a recording chamber on the stage of NiKon Eclipse Ti inverted microscope. NADH was excited at 350 nm (AT350/50X, Chroma) and fluorescence was recorded at 460 nm (ET460/50m and T400LP, Chroma). We normalized NADH level with FCCP as 0%, Rotenone-induced NADH change as 100%.

### Transmission electron microscopy and mitochondrial scoring

Hearts were fixed in 2.5 % gluteraldehyde in 0.1 M Na cacodylate buffer overnight at 4°C, washed 3 × 20 min in 0.1 M Na cacodylate buffer, pH 7.2, fixed in 4 % OsO_4_, washed in 0.1 M Na cacodylate buffer then dH_2_O, 2.5 % uranyl acetate, EtOH series to dehydrate then EtOH and Spurr’s with final solution 100% Spurr’s. Hearts were embedded in Spurr’s at 60°C for 24 – 48 h. Ultramicrotomy sections were cut at 90 nm and samples collected on 200 mesh for formvar grids for staining with uranyl and lead. Stained sections were examined with a Philips/FEI BioTwin CM120 Transmission Electron Microscope and digital images were collected with a Gatan Orius high-resolution cooled digital camera (2k×2k).

TEM images of mitochondria were scored by the following criteria: 0 = no detectable disruption in any mitochondria/field, 1 = cristae disrupted in one mitochondrion/field, 2 = disruption in >1 mitochondrion/field, 3 = ≥ 1 and < 50% ruptured mitochondria/field and 4 = ≥ 50% mitochondria ruptured/field. At least 10 images were scored per sample by a blinded technician and the mean score reported.

### Quantitative PCR

Total DNA was prepared using DNeasy Blood and Tissue Kit (Qiagen). Validated primers for Mitochondrial (CCCATTCCACTTCTGATTACC, ATGATAGTAGAGTTGAGTAGCG) and nuclear (GTACCCACCTGTCGTCC, GTCCACGAGACCAATGACTG) genes ^105^ were used for qPCR on CFX Connect™ Real-Time PCR Detection System (Bio-Rad). Mitochondrial to nuclear DNA ratios were quantified using the ΔΔCt method.

Total RNA was prepared using Trizol reagent. iScript™ Reverse Transcription Supermix (Bio-Rad) was used to generate cDNA from RNA. Validated PrimePCR primers (Bio-Rad) (Ccl2 qMmuCED0048300, Ccl3 qMmuCED0044190) and SsoAdvanced™ Universal SYBR^®^ Green Supermix (Bio-Rad) were used for qPCR on CFX Connect™ Real-Time PCR Detection System (Bio-Rad). Transcript levels were quantified by the ΔΔCt method.

### TUNEL

Cryosections (10μM) of ventricular tissue were fixed in 4% paraformaldehyde and stained with In-situ Cell Death Detection kit (Roche). Sections were imaged on a laser-scanning confocal microscope (LSM 510, Carl Zeiss). TUNEL positive and total nuclei were counted from 5 images per sample, and the averages reported.

### Fibrosis

Mouse hearts were formalin fixed and paraffinized. Heart sections were cut along the coronal plane at a thickness of 4 microns and mounted onto slides. The sections were kept at 60 degrees for 30 minutes and deparaffinized through xylene, 100% ethanol, 95% ethanol and then water. The slides were then stained using a Masson’s Trichrome Aniline Blue Stain Kit (Newcomer Supply, 9179B) and imaged at 20× with an Aperio Scanscope CS. The images were then analyzed with Aperio ImageScope, using a lower intensity threshold of 150 and a hue value of 0.66. Six left ventricular free wall images were analyzed per sample. The positivity percentages of the six images were then averaged and the average positivity values were reported.

### Liquid chromatography and mass spectrometry

Frozen extracted mitochondria were lysed by sonication in 300 μl of lysis buffer (8 M urea, 50 mM ammonium bicarbonate, 1X protease inhibitors (Complete mini EDTA-free mixture [Roche Applied Science], 1X phosphatase inhibitor mixture [PhosSTOP, Roche Applied Science]). After centrifugation (20,000 × g for 10 min at 4°C), the protein concentration of the supernatant was measured using the Bradford assay (BioRad) and ~150 μg of proteins for each condition were subjected to digestion. Protein reduction and alkylation were performed using a final concentration of 2 mM dithiothreitol and 4 mM iodoacetamide, respectively. Proteins were first digested for 4 h at 37°C with Lys-C (enzyme/substrate ratio 1:100). The second digestion was performed overnight at 37°C with trypsin (enzyme/substrate ratio 1:100) in 2 M Urea. The resulting peptides were chemically labeled using stable isotope dimethyl labeling as described previously ^106^. After protein digestion the mtCaMKII mitochondria were labeled as “Intermediate”, while the WT were labeled with “Light”. Sample were mixed in a 1:1 ratio and ~300 μg of the peptide mixtures were subjected to phosphopeptide enrichment using Ti^4+^-IMAC material as described previously ^107^. Briefly, the mixtures of labeled samples were dried to completion and reconstituted in 80% ACN, 6% trifluoroacetic acid (TFA) and loaded onto the Ti^4+^-IMAC columns. After washing with 50% ACN, 0.5% TFA, 200 mM NaCl and 50% ACN, 0.1% TFA consecutively, the phosphopeptides were eluted first with 10% ammonia and then with 80% ACN, 2% FA and were dried to completion in a vacuum centrifuge. After reconstitution in 10% FA, 5% dimethyl sulfoxide, the peptides were analyzed using nano flow reverse phase liquid chromatography on a Proxeon Easy-nLC 1000 (Thermo Scientific) coupled to an Orbitrap Elite (Thermo, San Jose, CA). Peptides were separated on an in-house made 50 cm column, 75 μm inner diameter packed with 1.8 μm C18 resin (Agilent Zorbax SB-C18) at a constant temperature of 40°C, connected to the mass spectrometer through a nanoelectrospray ion source. The injected peptides were first trapped with a double fritted trapping column (Dr Maisch Reprosil C18, 3 μm, 2 cm × 100 μm) at a pressure of 800 bar with 100% solvent A (0.1 % formic acid in water) before being chromatographically separated by a linear gradient of buffer B (0.1% formic acid in acetonitrile) from 7% up to 30% in 170 minutes at a flow rate of 150 nl/min. Nanospray was achieved with an in-house pulled and gold coated fused silica capillary (360 μm outer diameter, 20 μm inner diameter, 10 μm tip inner diameter) and an applied voltage of 1.7 kV. Full-scan MS spectra (from m/z 350 to 1500) were acquired in the Orbitrap with a resolution of 30,000. Up to ten most intense ions above the threshold of 500 counts were selected for fragmentation. For the fragmentation a decision tree method was used as described previously ^108^. The mass spectrometry proteomics data have been deposited to the ProteomeXchange Consortium via the PRIDE ^109^ partner repository with the dataset identifier PXD004631.

### LC-MS Data Analysis

For the raw data files recorded by the mass spectrometer, peak lists were generated using Proteome Discoverer (version 1.3, Thermo Scientific, Bremen, Germany) using a standardized workflow. Peak list was searched against a Swiss-Prot database (version 2.3.02, taxonomy Mus musculus, 32402 protein entries) supplemented with frequently observed contaminants, using Mascot (version 2.3.02 Matrix Science, London, UK). The database search was performed by using the following parameters: a mass tolerance of 50 ppm for the precursor masses and ±0.6 Da for CID/ETD fragment ions. Enzyme specificity was set to Trypsin with 2 missed cleavages allowed. Carbarmidomethylation of cysteines was set as fixed modification, oxidation of methionine, dimethyl labeling (L, I) of lysine residues and N termini, and phosphorylation (S, T, Y) were used as variable modifications. Percolator was used to filter the PSMs for <1% false discovery-rate. Phosphorylation sites were localized by applying phosphoRS (pRS) (v2.0) ^110^. Double dimethyl labeling was used as quantification method ^111^, with a mass precision of 2 ppm for consecutive precursor mass scans. A retention time tolerance of 0.5 min was used to account for the potential retention time shifts due to deuterium. To further filter for high quality data we used the following parameters: high confidence peptide spectrum matches, minimal Mascot score of 20, minimal peptide length of 6 only unique rank 1 peptide and the search rank 1 peptide. The phosphopeptides that showed an on/off situation in the mtCaMKII or WT were manually quantified by giving them an arbitrary value of 100 or 0.01 for extreme up- or down-regulation, which corresponds to the maximum allowed fold change in the used Proteome Discoverer settings.

### Modeling and simulation

A three compartment computational model was constructed to include mitochondrial matrix, intermembrane space (IMS), and cytosol compartments. The compartments were separated by inner mitochondrial membrane (IMM), and outer mitochondrial membrane (OMM). The computational model consists of 48 nonlinear ordinary differential equations. The mitochondrial processes include oxidative phosphorylation, TCA cycles and transporters across the mitochondrial inner membrane ^63, 64^. The mitochondrial creatine kinase in the intermembrane space (IMS) and in the cytosol were modeled as random sequential bi-bi enzyme reactions and creatine, creatine phosphate, ATP and ADP were diffused between IMS and cytosol ^112^. The equations were solved by ODE solver ode15s in MATLAB 2016a (The MathWorks, Natick, MA). The computational model of mitochondrial and cellular CaMKII and creatine kinase system was constructed to simulate over-expressed mitochondrial CaMKII condition and mitochondrial creatine kinase (CK). The mitochondrial CaMKII overexpressing (mtCaMKII) condition was simulated by decreasing complex I activity to 45% of control, decreasing mitochondrial CK to 80% of control, and increasing complex II (130%) and TCA cycle enzymes activities (isocitrate dehydrogenase (IDH) 120%,α-ketoglutarate dehydrogenase(αKGDH) 120%, fumarate hydratase (FH) 120%, malate dehydrogenase (MDH) 20% more) according to the experimental measurements of these enzyme activities. Mitochondrial CK over-expression was simulated by increasing mitochondrial CK concentration to 120% of control. ATP hydrolysis during cardiac systole was simulated using a pulsatile function. Simulation codes are available at the URL: https://gitlab.com/MitoModel/mtCaMKII.git.

### Statistical Analysis

Statistical analyses were performed using Graph Pad Prism 7 software. Sample size and information about statistical tests are reported in the figure legends. Data are presented as mean ± SEM. Pairwise comparisons were performed using a two-tailed t test. For experiments with more than 2 groups, data was analyzed by 1 way ANOVA followed by Tukey’s post-hoc multiple comparisons test.

## Data Availability

The data that support the findings of this study are available from the corresponding authors upon reasonable request.

## Acknowledgements

This work was supported by the NIH (R35 HL140034 to M.E.A. and HL63030 and HL61912 to R.G.W.), an American Heart Association Collaborative Sciences Award (17CSA33610107 to M.E.A.), and the Netherlands Organization for Scientific Research (NWO) through funding of the large-scale proteomics facility *Proteins@Work* (project 184.032.201) embedded in the Netherlands Proteomics Centre (E.C. and A.J.R.H.). We thank Jinying Yang for animal model maintenance; Marwan Mustafa, Djahida Bedja, Michelle Leppo, and the Cardiovascular Physiology and Surgery Core at Johns Hopkins School of Medicine for technical assistance; Shawn Roach and Teresa Ruggle for graphic design and figure preparation.

## Author Contributions

Conceptualization, E.D.L, M.E.A.; Software, A.W.; Methodology, Y. Wang; Validation, E.D.L.; Formal Analysis, E.D.L., A.W., A.G., J.M.G, N.R.W., K.R.M., P.U.; Investigation E.D.L, J.M.G., N.R.W., Y.Wu, A.W., A.G., E.C., M.A.J., A.S., O.E.R.G, K.R.M., P.U.; Data Curation, E.D.L.; Writing-Original Draft, E.D.L. and M.E.A.; Writing-Review & Editing, E.D.L., M.E.A, R.G.W., A.W.; Supervision M.E.A., A.J.R.H., R.G.W.; Project Administration, E.D.L.; Funding Acquisition, M.E.A, A.J.R.H., and R.G.W.

## Competing Interests

M.E.A. is a named inventor on patents claiming to treat heart disease by CaMKII inhibition. He is a founder and advisor to Allosteros Therapeutics. At present no income derives from these activities.

---

**Figure S1.**
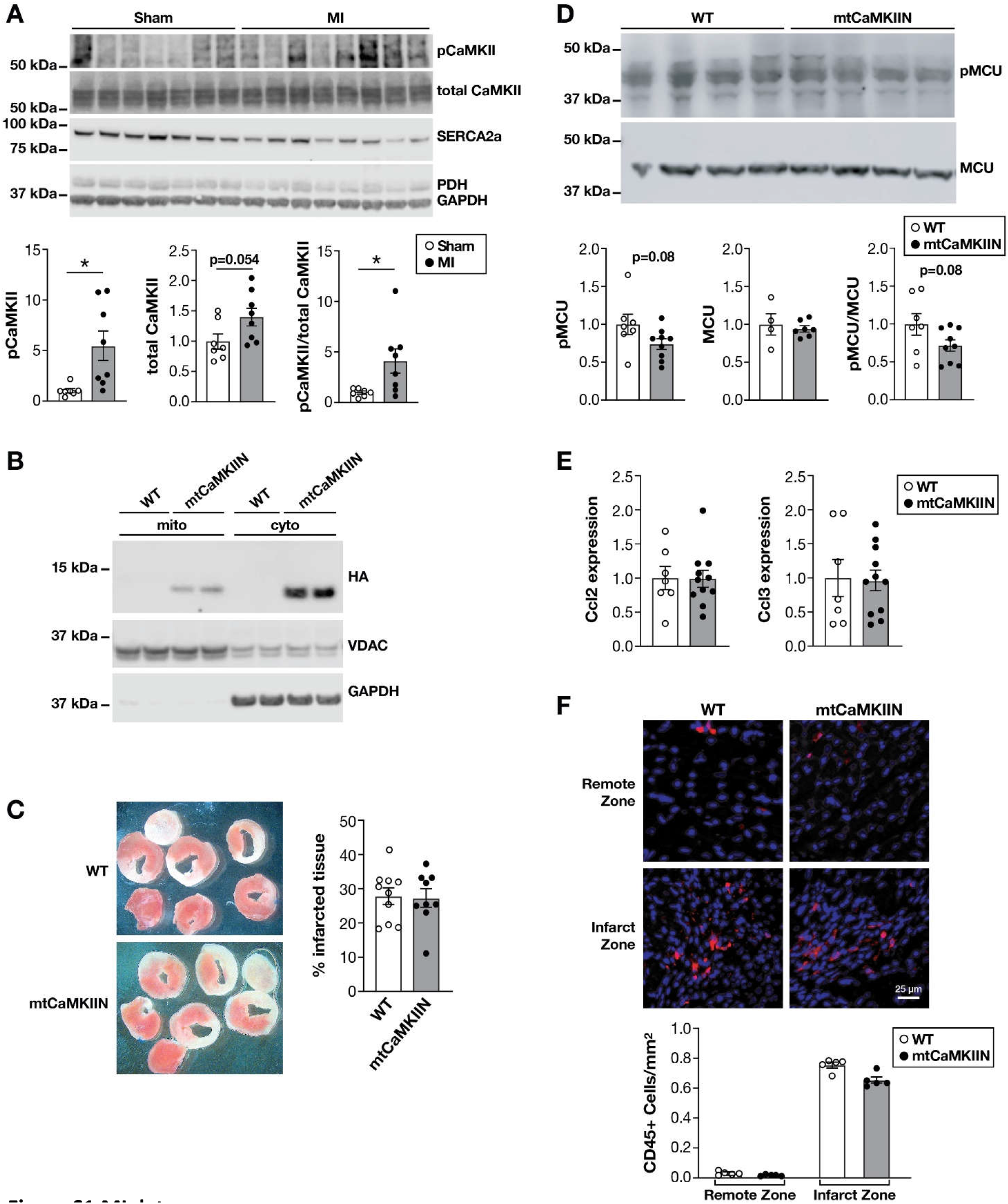
MI data. (A) Western blot of cytosolic lysates and summary data for phosphorylated CaMKII and total CaMKII normalized to coomassie staining for sham (n=7) and MI (n=8) mitochondria. Blots for cellular compartment markers also included: SERCA2a (SR membrane), PDH (mitochondrial matrix), GAPDH (cytosol). (B) Western blot for HA-CaMKIIN, VDAC1 and GAPDH in cytoplasm and mitochondria fractions from WT (n=2) and mtCaMKIIN hearts (n=2). (C) Representative images and quantification of of TTC staining in WT and mtCaMKIIN hearts 24 hours after MI surgery. (D) Representative western blot and summary data of pMCU and MCU normalized to coomassie in mitochondrial lysates from WT (n=7 and mtCaMKIIN (n=9) 1 week after MI. (E) qPCR quantification of Ccl2 and Ccl3 mRNA normalized to Hprt in WT (n=7) and mtCaMKIIN (n=11) hearts 1 week after MI. (F) Representative immunofluorescence images and summary data for CD45+ cells in heart sections of WT (n=5 hearts) and mtCaMKIIN (n=5 hearts) hearts 1 week after MI. (Data are represented as mean α SEM, significance was determined using two-tailed t test, (*P<0.05)

**Figure S2.**
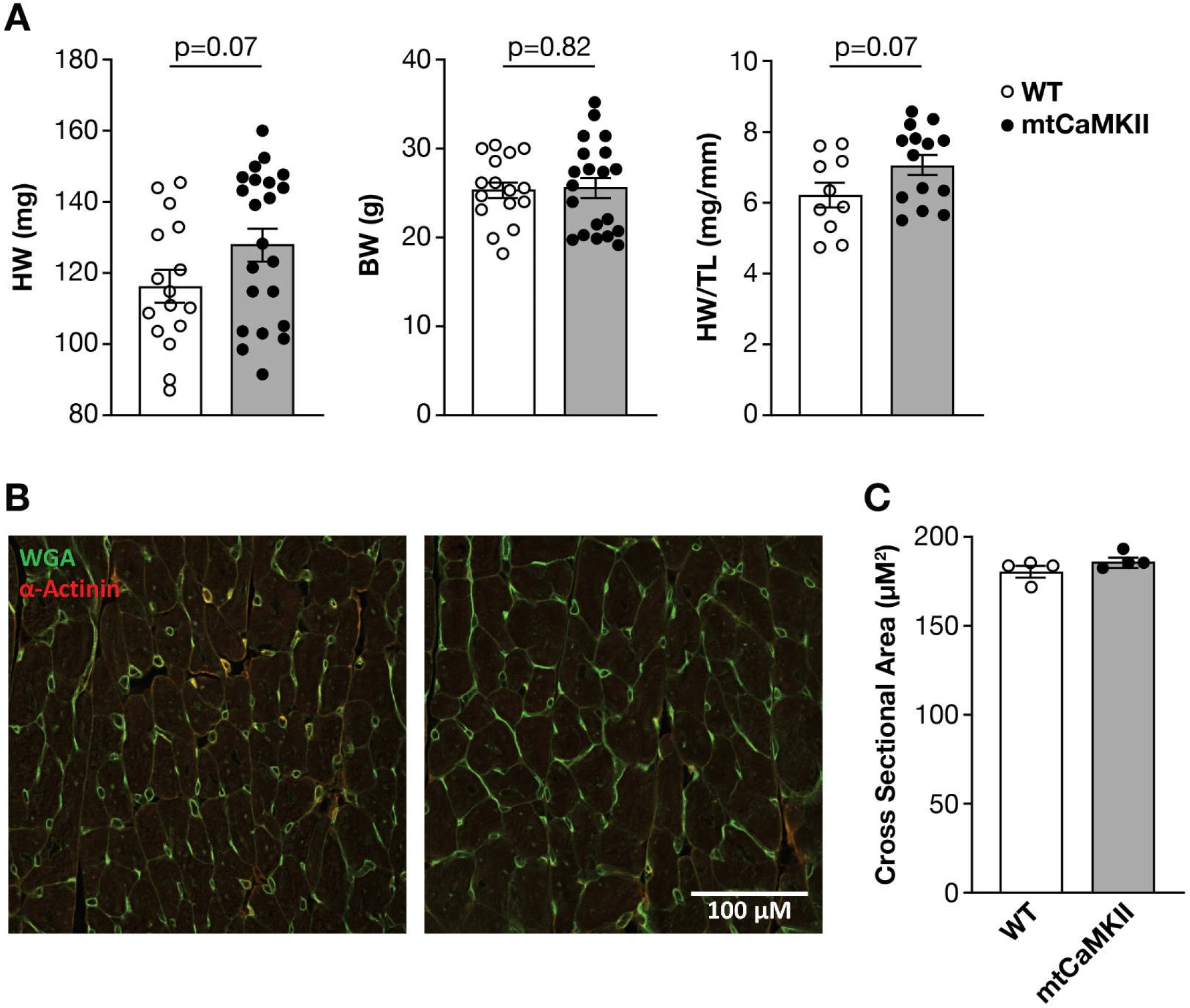
mtCaMKII mouse data. (A) Summary data for HW, BW and HW/TL measurements for WT (n=10-16) and mtCaMKII (n=14-22) mice. (B) Representative images and (C) summary data for quantification of cardiomyocyte cross sectional area in WT (n=4 hearts) and mtCaMKII (n=5 hearts) heart sections. (Data are represented as mean ± SEM, significance was determined using two-tailed t test.)

**Figure S3.**
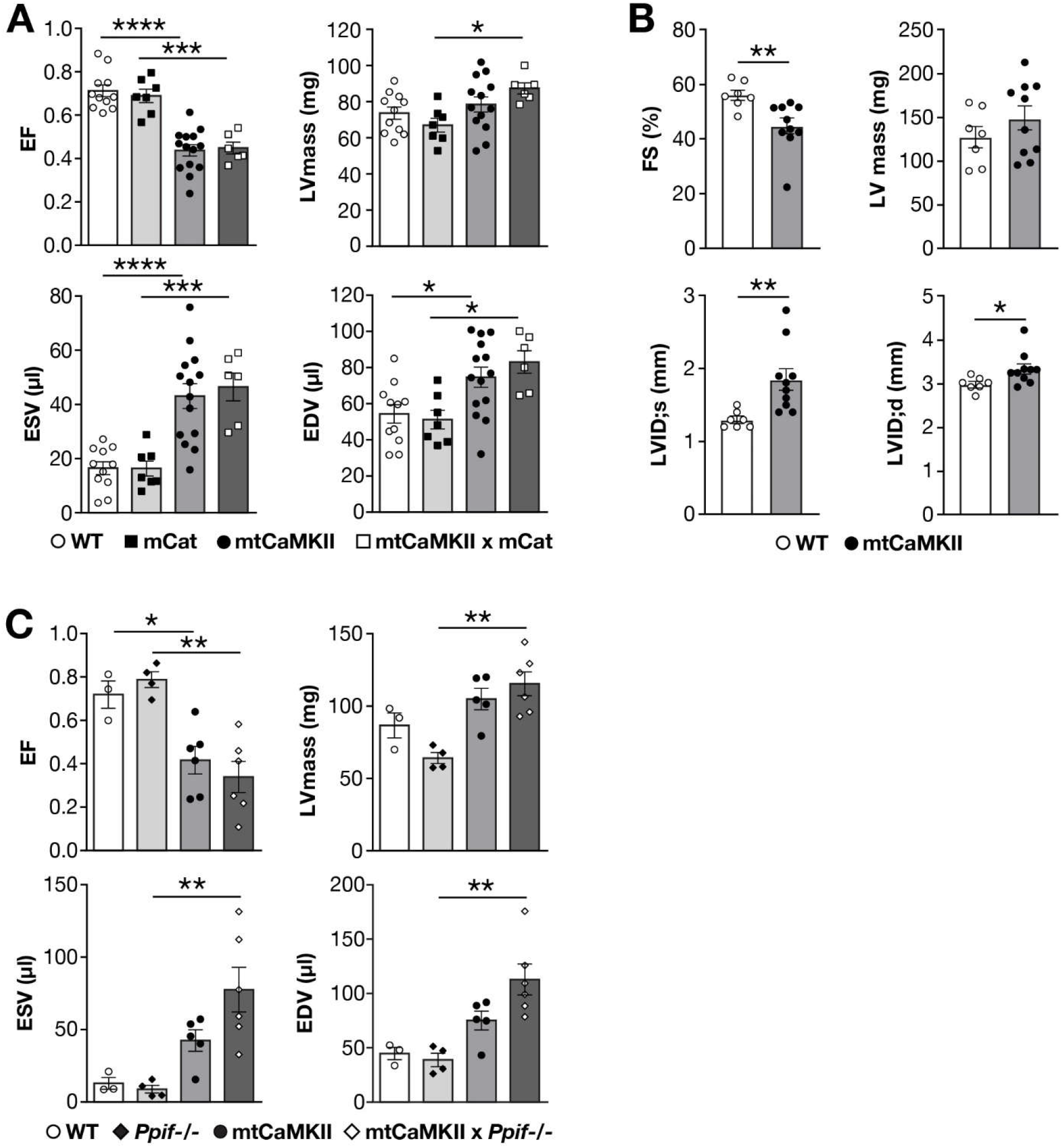
Genetic crosses do not rescue mtCaMKII dilated heart phenotype. (A) Summary data from echocardiographic measurements in WT (n=11), mitochondrial-targeted catalase over-expressing (mCat, n=7), mtCaMKII (n=14), and mtCaMKII x mCat interbred (n=6) mice, (B) WT CD1 (n=7) and mtCaMKII CD1 (n=10), and (C) WT (n=4), *Ppif*^−/−^ (n=4), mtCaMKII (n=6), and mtCaMKII x *Ppif*^−/−^ interbred (n=6) mice. (Data are represented as mean ± SEM, significance was determined using two-tailed t test or 1 way AVOVA with Tukey’s multiple comparison’s test, ****P<0.0001, ***P<0.001, **P<0.01, *P<0.05)

**Figure S4.**
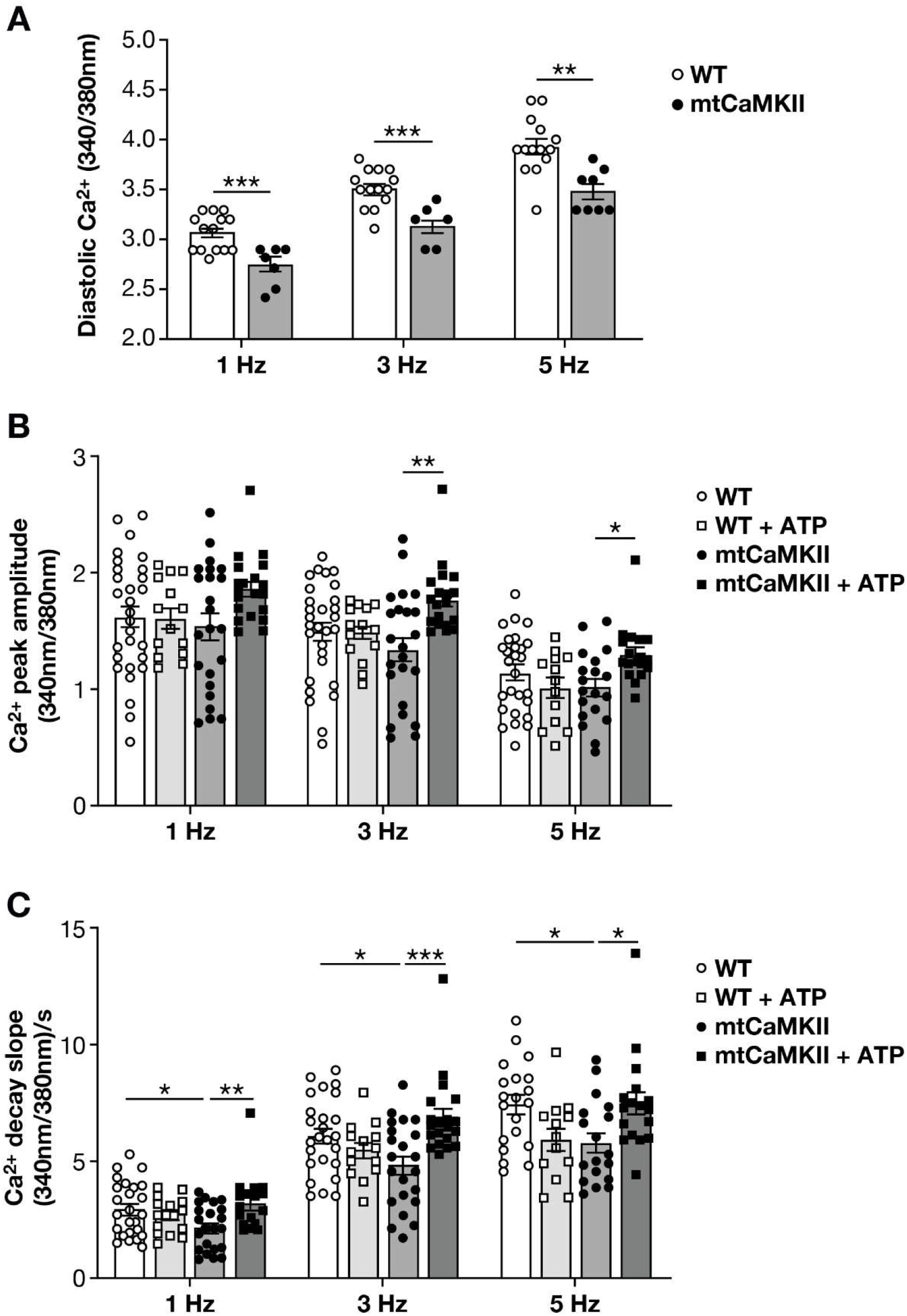
[Ca^2+^] in myocytes. (A)Summary data for diastolic [Ca^2+^] measurements made with Fura-2–loaded ventricular myocytes isolated from WT (14 cells from 2 hearts) and mtCaMKII (8 cells from 2 hearts) hearts and field stimulated (1, 3, and 5 Hz). Myocyte cytoplasm was not perfused with a CaMKII inhibitory peptide (AIP) as in Figs 3E-F and 4H. Summary data for (B) Ca^2+^ peak amplitude and (C) decay slope from isolated ventricular myocytes stimulated at 1, 3, or 5 Hz (n=12-19 ventricular myocytes isolated from 2-3 hearts/group). (Data are represented as mean ± SEM, significance was determined using a two-tailed t test, or 1 way AVOVA with Tukey’s multiple comparison’s test ***P<0.001, **P<0.01, *P<0.05)

**Figure S5.**
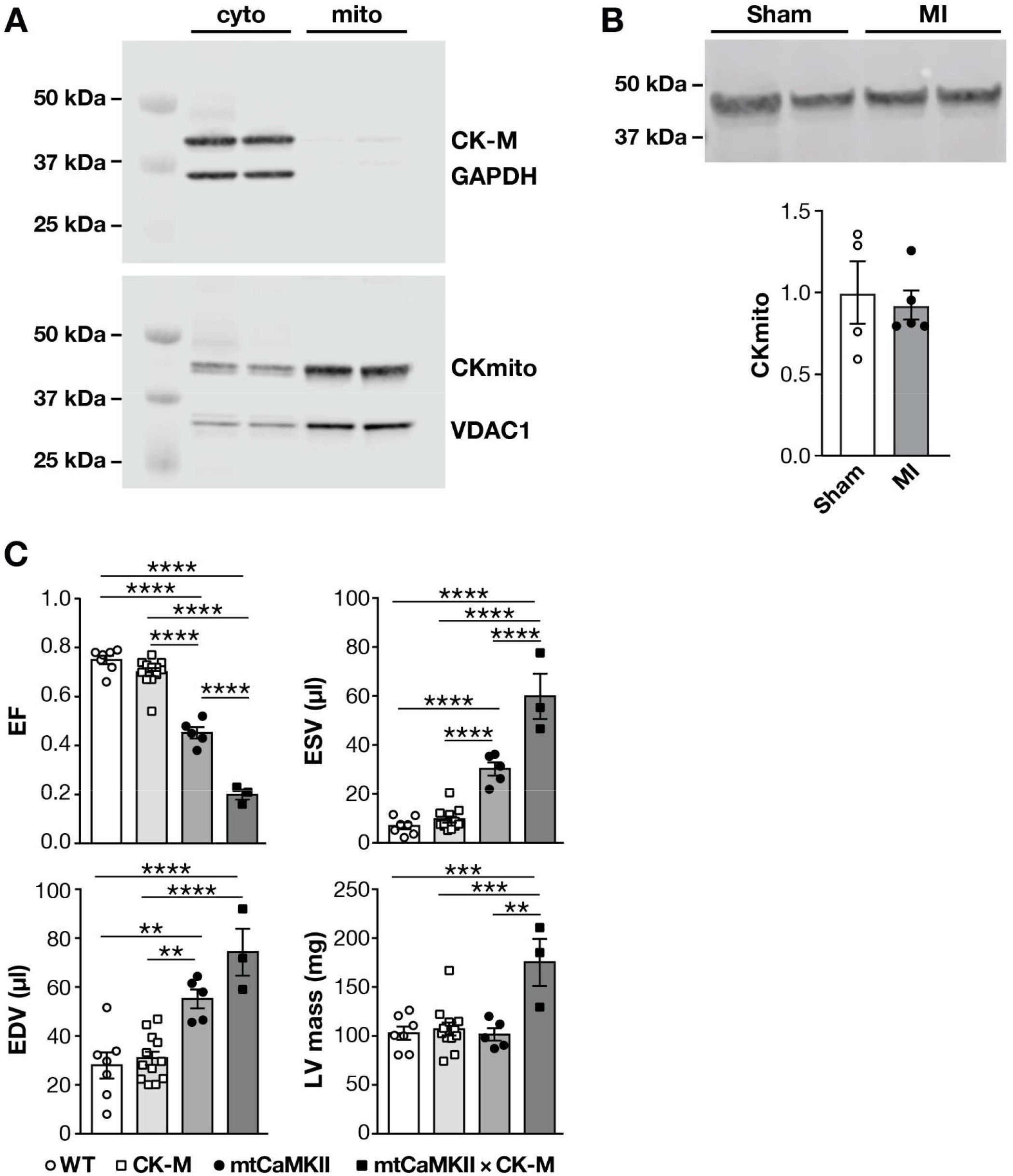
CK expression in transgenic and MI hearts. (A) Western blot for CKmito and CK-M of cytosolic and mitochondrial fractions from WT mouse hearts. (B) Representative western blot and summary data for Ckmito in sham (n=4) and MI (n=5) hearts. (C) Summarized echocardiographic measurements from WT (n=7), CK-M (n=12), mtCaMKII (n=5), and mtCaMKII x CK-M (n=3) mice. (Data are represented as mean ± SEM, significance was determined using 1 way AVOVA with Tukey’s multiple comparison’s test, ****P<0.0001, ***P<0.001, **P<0.01)

**Figure S6.**
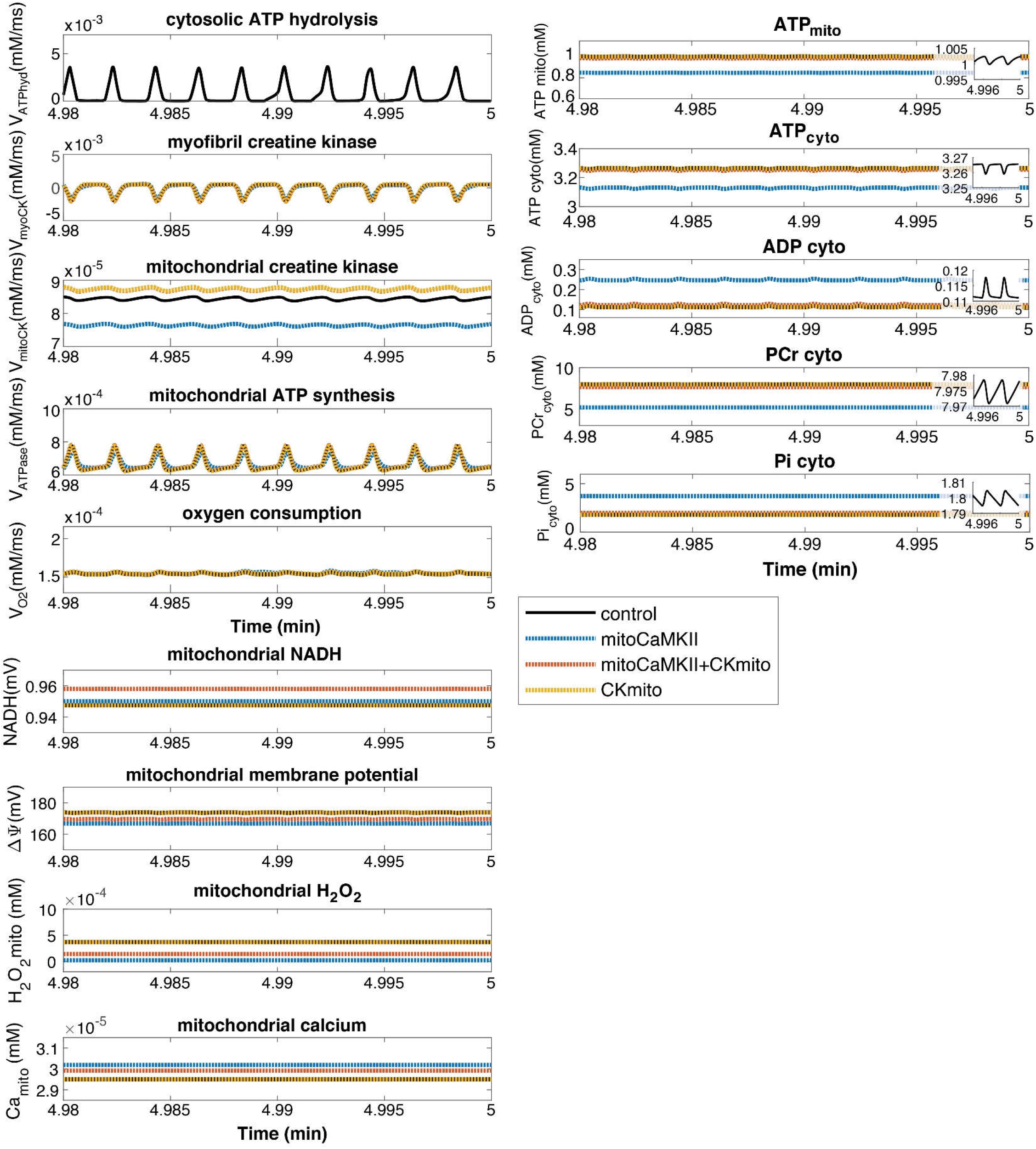
Computational Model Outputs. ATP hydrolysis was simulated using a pulsatile function to simulate cardiac contraction. Mitochondrial creatine kinase flux, myocardial creatine kinase flux, mitochondrial ATP synthesis rate, and oxygen consumption as well as mitochondrial ATP, cytosolic ATP, ADP, PCr, Pi levels, and mitochondrial NADH, membrane potential, H_2_O_2_ and Ca^2+^ were compared in WT, CKmito, mtCaMKII, and mtCaMKII x CKmito interbred conditions after 5 minutes of simulation. Insets in right panels provide zoomed in scale to show small oscillations.

**Table S1.** CaMKII has diverse mitochondrial targets. Proteins identified by LC/MS analysis to have increased phosphorylation in mtCaMKII compared to WT mitochondria. Peptides were considered to be significantly more phosphorylated with a fold change of ≥ 2 and P<0.01. Proteins were classified by function.

| Pathway |  | Phosphoproteins |  |  |
| --- | --- | --- | --- | --- |
| Metabolism | TCA Cycle | ACON<br>ODO1<br>ODPX | CISY<br>ODO2<br>SUCB1 | MDHM<br>ODPA |
|  | Electron transport/<br>Oxidative Phosphorylation | ATP5H (CV)<br>UQCC2 (CIII) | COX41 (CIV) | NDUFB11 (CI) |
|  | Fatty Acid Oxidation | ACADV<br>ETF A | ACSL1 | ECI1 |
|  | Other Metabolic | AATM<br>GCSH | ADAS<br>KCRS | ARK72<br>MUTA |
| Transcription/Translation |  | EFTS<br>RM01<br>SLIRP | NUCKS<br>RT02<br>SYIM | PERM1<br>RT23 |
| Oxidative Stress |  | MSRB2 | PRDX3 | PRDX5 |
| Import |  | TIM44 | TOM70 |  |
| Fission/Fusion |  | MFR1L |  |  |
| Unknown |  | CQ059 | OCAD1 |  |

**Table S2.** Parameters from experimental data used for computation modeling. Summary of the parameters used in the computational model that were taken from the experimental data for WT, CKmito, mtCaMKII, and mtCaMKII x CKmito.

| Parameters | WT | CKmito | mtCaMKII | mtCaMKII x CKmito | Source |
| --- | --- | --- | --- | --- | --- |
| **Complex I expression/activity** | 100% | 100% | 45% | 70% | Fig 5E and H |
| **CKmito expression/activity** | 100% | 120% | 80% | 120% | Fig 4E and F |
| **CKmito forward rate constant** | 100% | 120% | 80% | 120% |  |
| **Cytosolic ATP hydrolysis** | 100% | 100% | 100% | 100% |  |
| **TCA cycle enzyme activity (IDH,aKGH, FH, MDH)** | 100% | 100% | 120% | 120% | Fig 6B |
| **Complex II expression/activity** | 100% | 100% | 130% | 130% | Fig 5E |

